# GDF5 as a Multimodal Protector of the Motor Unit in Amyotrophic Lateral Sclerosis

**DOI:** 10.64898/2026.06.30.735251

**Authors:** Aly Bourguiba, Sonia Pezet, Massiré Traoré, Christel Gentil, Thibaut Marais, Aurélie Fail, Maxime Gelin, Julien Mésseant, Pierre Meunier, Sofia Benkhelifa-Ziyyat, Zoheir Guesmia, Bruno Cadot, Antonio Musaro, Gabriella Dobrowolny, Julianne Perronnet, Sestina Falcone, Piera Smeriglio, France Pietri-Rouxel

## Abstract

Amyotrophic lateral sclerosis is characterized by the progressive dismantling of the motor unit. While “dying back” hypothesis suggests that peripheral neuromuscular dysfunction precedes motor neuron loss, the molecular mechanisms limiting endogenous compensatory responses remain poorly understood. We longitudinally examined neuromuscular decline and GDF5-SMAD1/5/8 signaling in SOD1^G93A^ mice. Our findings revealed a translational checkpoint linked to the lncRNA *Myoparr* that suppresses GDF5 production at symptom onset. To overcome this deficit, we delivered AAV9-GDF5 at the symptomatic stage. GDF5 supplementation restored SMAD signaling balance, shifting the motor unit from a pro-atrophic TGF-β-SMAD2/3 toward a pro-myogenic SMAD1/5 profile. Treatment preserved muscle mass, reduced mitochondrial reactive oxygen species, and maintained neuromuscular junction integrity, including peri-synaptic glial support. GDF5 also promoted molecular recovery of spinal MNs by enhancing homeostatic marker expression. Together, these findings identify GDF5 as a multimodal stabilizer of the motor unit and highlight its potential as therapeutic target in combinatorial strategies aimed at coupling motor unit stabilization with central neuroprotective interventions.

## Introduction

Amyotrophic Lateral Sclerosis (ALS) is the most common adult-onset motor neuron (MN) disease, with a lifetime risk approximately 1 in 400(1,2). This fatal disease is characterized by degeneration of both lower MNs, in the brainstem and spinal cord, and upper MNs in the motor cortex(3) and as a consequence leads to severe muscle atrophy. Two forms have been described for ALS: sporadic (sALS), 90-95% of cases with different etiologies, or familial (fALS) less frequent, 5-10% of cases, and having genetic origins^4,5^. To date, no definitive cure for ALS exists. Although a limited number of pharmacological treatments has been approved, they offer only modest benefits in slowing disease progression(6). Consequently, gene therapy has emerged as a promising strategy for targeting fALS types, like the one associated to mutation in *superoxide dismutase1* (SOD1) gene(4,7). *SOD*1 mutation is one of the most common causes of fALS inducing the misfolding of the coded protein which leads to its aggregation in endoplasmic reticulum (ER) of MNs. While this process impairs SOD1 role to catalyze the elimination of free radicals and reactive oxygen species (ROS), the pathophysiology is primarily driven by a toxic gain of function stemming from the presence of the misfolded aggregates(8). While significant pharmacological progress has been made, most notably with the approval of Tofersen, an antisense oligonucleotide targeting *SOD1* mRNA, its clinical application is strictly limited to patient harboring mutations in this specific gene (9). However, given the predominance of sporadic cases, the development of therapeutic approaches effective for both, fALS and sALS, remains a critical unmet need.

Beyond the known primary damage affecting MN, ALS is increasingly recognized as a systemic disease characterized by the dysregulation of various muscle signaling pathways. Notably, recent studies revealed an upregulation and aberrant activation of the transforming growth factor beta (TGF-β) signaling pathway in both skeletal muscle and spinal cord of ALS patients(10). In muscle, TGF-β induces the expression of E3 ubiquitin ligases such as MuRF1, MUSA1, and Atrogin-1, promoting Ubiquitin/Proteasome-mediated (UP) protein degradation and atrophy. This catabolic action is normally counterbalanced by the bone morphogenetic protein (BMP) pathway, which supports muscle growth and maintenance, suggesting that disruption of this equilibrium contributes to ALS-related muscle wasting(11,12). In this context, Growth Differentiation Factor 5 (GDF5), also known as BMP 14, a BMP family member that activates SMAD1/5/8 (fusion of *Caenorhabditis elegans* Sma and the *Drosophila* Mad, or mothers against decapentaplegic) signaling, was described to play key roles in the neuromuscular system(11,13–15). GDF5 has been described to support neurites growth and neuron survival *in vitro*(16,17). Widely, its signaling pathway SMAD 1/5/8 has been shown to decrease neuronal excitotoxicity(18). GDF5 is also considered a muscle trophic factor. Indeed, GDF5 has been demonstrated as required for skeletal muscle re-innervation after nerve crush and limiting denervation related muscle atrophy(11,15). In addition, we were pioneers in discovering that GDF5 supplementation in aged mice promotes muscle hypertrophy and control proteostasis by increasing protein synthesis and decreasing protein degradation depending on ubiquitin ligases(13,14).

Given the dual neurotrophic and myotrophic described effects of GDF5, we therefore hypothesized that GDF5-based treatment could preserve innervation, muscle mass and function, having thus a potential beneficial impact on the disease progression of both, fALS and sALS addressing the whole motor unit.

In this study, using the high-copy SOD1^G93A^ transgenic mouse model of ALS, we first showed abnormal expression of GDF5 in muscle of these mice and that the balance between BMP and TGF-β was disrupted. In addition, carrying out bulk RNA sequencing (RNA-Seq) revealed significant alteration in stress responses, particularly in apoptosis, oxidative stress and unfolded protein response in muscle of the SOD1^G93A^ mice. Secondly, we demonstrated that GDF5 overexpression (GDF5 OE) mediated by associated adenovirus (AAV) administrated through systemic route, has a beneficial effect on the ALS pathophysiology in preserving muscle mass, structure and function. Comparing gene expression profiles of treated and untreated SOD1^G93A^ mouse muscle by RNA-Seq, showed that GDF5 OE increased oxidoreductase activity and muscle hypertrophy pathway. Furthermore, the analysis of neuromuscular junctions (NMJ) revealed an improvement in their morphology and in innervation status.

Overall, this study paves the way for GDF5-based treatment that could potentially be applied to all forms of ALS, with the aim of improving both muscle and motor neuron pathophysiology and thus slowing the overall progression of ALS.

## Results

### SOD1^G93A^ mice display progressive muscle denervation

The SOD1^G93A^ mouse model is a well-established murine model of ALS that overexpresses the human mutated gene of *SOD1* gene and recapitulates the major stages of disease progression(19). Longitudinal monitoring of disease progression showed that mice remain clinically asymptomatic during early life, develop the first motor symptoms around postnatal day 80 (P80), corresponding to the early symptomatic stage, and become severely paralyzed by approximately P120 (late symptomatic stage), ultimately reaching end-stage disease shortly thereafter (19) (Fig 1A). To further characterize neuromuscular features at the different stages, we analyzed muscle mass, neuromuscular transmission as well as NMJ remodeling markers. At symptom onset, P80, SOD1^G93A^ mice presented Tibialis anterior (TA) muscle atrophy compared to Wild Type mice (WT), which progressively worsened during the late symptomatic phase up to P120 (Fig 1B). Neuromuscular transmission, evaluated by electroneuromyography (ENMG) before (P60) and at the mid-symptomatic period (P90-P100), revealed a decrement of the compound muscle action potential (CMAP) compared to WT mice (Fig 1C).

**Figure 1:**
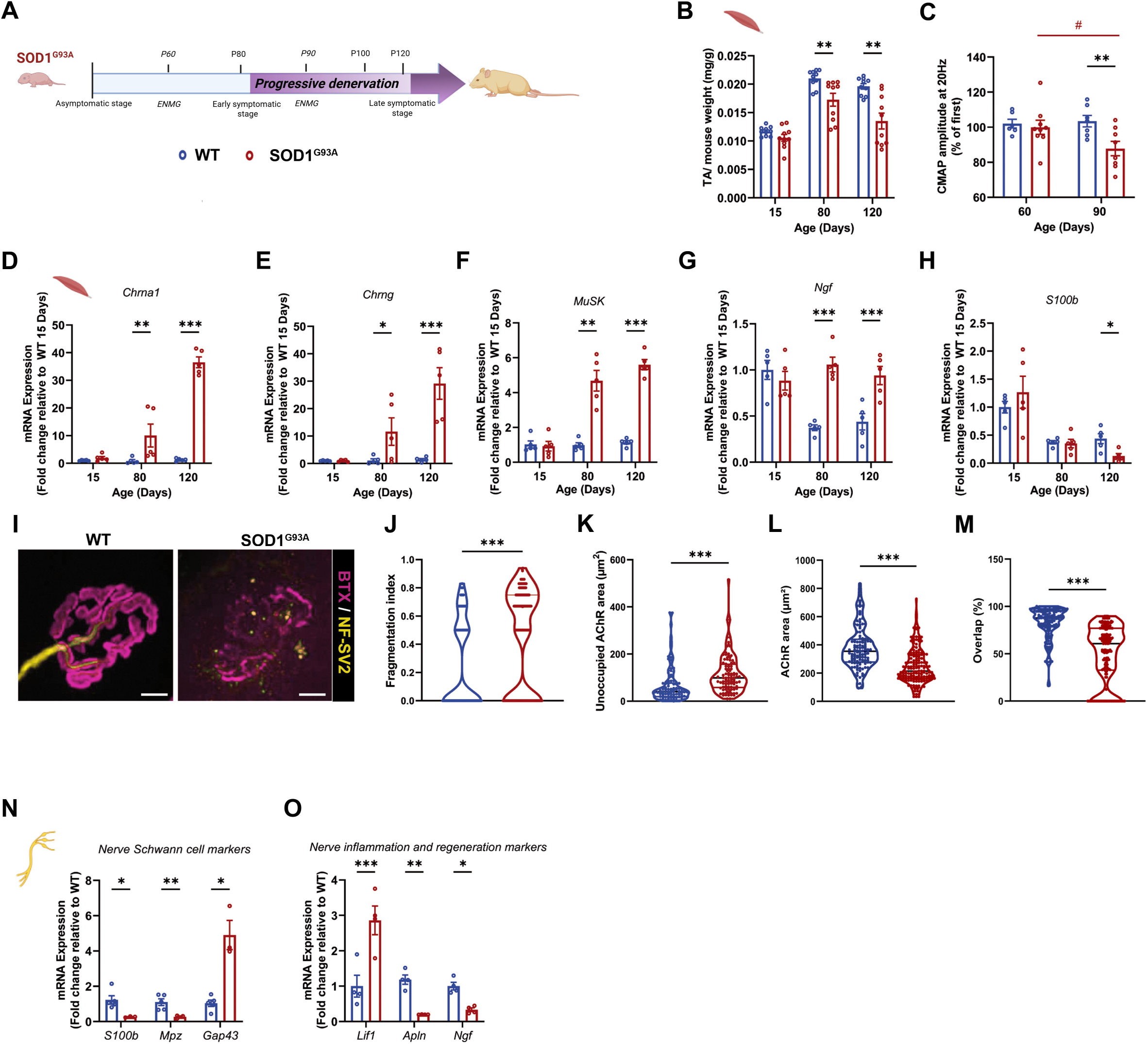
SOD1^G93A^ mice display progressive muscle denervation. **A:** Schematic representation of the natural history of the disease in the SOD1^G93A^ mouse model highlighting asymptomatic (P15), early symptomatic (P80), mid symptomatic (P90-P100)and late symptomatic (P120) stages **B:** Quantification of Tibialis anterior (TA) mass normalized on mouse weight in percentage (mg/g) in WT (blue) and SOD1^G93A^ (red) mice at P15, P80 and P120 (N=10) **C:** Electroneuromyography (ENMG) analysis showing the decrement of the compound muscle action potential (CMAP) in WT (blue) and SOD1^G93A^ (red) TA at P60 and P90 (N= 6-9) **D-H:** RT-qPCR analysis of NMJ remodeling markers *Chrna1* (D), *Chrng* (E), *Musk* (F), *Ngf* (G) and *S100b* (H) expression in TA muscle at P15, P80 and P120 (N= 5) **I:** Representative images of NMJ morphology from WT and SOD1^G93A^ mice at P120 analyzed using NMJ morph. Presynaptic and post synaptic region are stained with NF200 +SV2 (Yellow) and α-bungarotoxin (Magenta), scale bar 10 µm **J-M:** Quantification of NMJ hallmarks including: fragmentation index (J), unoccupied AChR area (K), total AChR area (L) and overlap percentage between pre- and post-synaptic component (M) (N= 124-166) **N:** RT-qPCR of glial support axonal inflammation/regeneration markers *S100b, Mpz, Gap43,* in the sciatic nerve at mid symptomatic stage (N= 4-5) **O:** RT-qPCR of axonal inflammation/regeneration markers *Lif, Apln* and *Ngf* in the sciatic nerve at mid symptomatic stage (N= 4-5) Data are presented as means ± s.e.m. P-values were calculated by Two-way ANOVA followed by Fisher LSD (B, C,D, E, F, G, H, N,O) ; Mann Whitney (J,K,L,M)

To further characterize denervation during disease progression, we quantified the expression of genes associated with NMJ remodeling, including the acetylcholine receptor α1 subunit (*Chrna1*), the γ subunit (*Chrng*), and MuSK (muscle-specific kinase), all of which are induced following denervation (14,20). We also measured the mRNA level of the *nerve growth factor* (*Ngf)*, a neurotrophin overexpressed in ALS muscle(21). The expression of all these markers was significantly increased from the early symptomatic stage (P80) and remained elevated until the late symptomatic stage (P120) (Fig 1D-G). Knowing that the integrity of the NMJ is linked to the abundance of terminal Schwann cells (SCs)(22), we quantified the transcript level of their marker, *S100b*(14,20) and we observed its decrease at the late symptomatic stage (P120) marking an alteration of these cells (Fig 1H). To corroborate muscle denervation and the variation of synaptic remodeling markers at P120, we analyzed NMJ morphology using an ImageJ-based workflow (Fig 1I)(14,23). We found an increase of the NMJ fragmentation index (Fig 1J), of the unoccupied AChR area Fig 1 K) in muscle of SOD1^G93A^ compared to WT mice. In parallel, a decrease in AChR area and in the percentage of overlap between pre- and postsynaptic regions was detected in muscles of ALS mice (Fig 1 L-M). Collectively, these data confirm that the neuromuscular system of SOD1^G93A^ mice undergoes profound and progressive disruption resulting from a combination of functional, molecular and structural defects.

To further describe the molecular state of the sciatic nerve during the symptomatic phase of the SOD1^G93A^ mice, we investigated the expression of markers associated with glial support and axonal regeneration at mid-symptomatic stage. The data showed a significant downregulation of the SCs and myelin-associated marker *S100b* and *Mpz* (Myelin protein zero), reflecting a progressive failure of the peripheral glial environment (Fig 1N). Concurrently, we detected an induction of *Gap43* (Growth-associated protein 43), a feature of axonal growth and regeneration attempts(24,25), alongside a significant increase in the neuropoietic cytokine *Lif* (Leukemia inhibitory factor)(26) (Fig 1O). Furthermore, this endogenous regenerative response was associated with a significant reduction in key neurotrophic factors, such as *Apln* (Apelin) and *Ngf* (27).

Altogether, these results define a precise kinetics of the neuromuscular degradation in SOD1^G93A^ model. We demonstrated that, in muscle of these mice, molecular remodeling of the NMJ and functional CMAP decrement occur as early as the first clinical signs preceding the structural collapse of the synaptic architecture and the massive atrophy observed at terminal stage (P120). In addition, the molecular profiling of the sciatic nerve highlighted a transition at mid-symptomatic stage (P90-P100), when an endogenous regenerative attempt is rapidly overwhelmed by the loss of glial support and neurotrophic supply. This sequence of events reveals a progressive and sustained collapse of the motor unit homeostasis throughout ALS progression.

### The GDF5-SMAD1/5/8 signaling pathway is impaired along the neuromuscular axis

The observation that endogenous repair mechanisms fail to maintain neuromuscular integrity (Fig1) suggests that key compensatory signaling pathways are either insufficient or inhibited. Among proteins involved in the compensatory response, GDF5 is a key actor driving a protective response that limits muscle atrophy and stabilizes synapses following denervation(11,15). We therefore sought to determine whether this endogenous GDF5-mediated response was effectively implemented or compromised as the disease progressed in SOD1^G93A^ mice. We observed a progressive upregulation of *Gdf5* transcript in TA muscle, increasing at P80 and intensifying by P120 in muscles of SOD1^G93A^ compared to WT mice (Fig 2A). At P15, we observed high levels of GDF5 protein in muscle, consistent with an active endogenous response. In addition, transcript levels of the extracellular BMP antagonists *Noggin* and *Follistatin* were not different from those observed in WT mice at P15 (Fig S1A). These findings argue against extracellular sequestration of GDF5 as the cause of the lack of signaling activity and instead suggest the existence of an intracellular blockade preventing pathway activation despite elevated GDF5 protein levels. Surprisingly, GDF5 protein became undetectable at P80, despite persistently elevated *Gdf5* transcript levels (Fig 2B-C). The discrepancy between mRNA abundance and protein expression strongly suggested the existence of post-transcriptional or translational regulatory mechanisms. To investigate this possibility, we quantified the expression of the long non-coding RNA (lncRNA) *Myoparr*, previously identified as a translational inhibitor of GDF5. Consistent with this hypothesis, *Myoparr* expression was high at P15 in both WT and SOD1^G93A^ muscles. However, whereas its expression progressively declined in WT muscles with age, it remained markedly elevated in SOD1^G93A^ muscles at P80 (Fig 2D). The persistence of high *Myoparr* levels at this stage provided a plausible explanation for the absence of detectable GDF5 protein despite sustained *Gdf5* transcription. At the late symptomatic stage (P120), GDF5 protein levels increased sharply compared with WT muscles (Fig 2B). This rise was accompanied by significant upregulation of the downstream target genes *Id1* and *Id2*, as well as *Follistatin* (Fig S1A-B), indicating activation of GDF5 signaling. Notably, this restoration of GDF5 protein expression occurred despite the continued presence of high *Myoparr* levels. This observation suggests that the translational repression exerted by *Myoparr* can be overcome when *Gdf5* transcript abundance reaches a critical threshold, implying that *Myoparr*-mediated inhibition is saturable rather than absolute.

**Figure 2:**
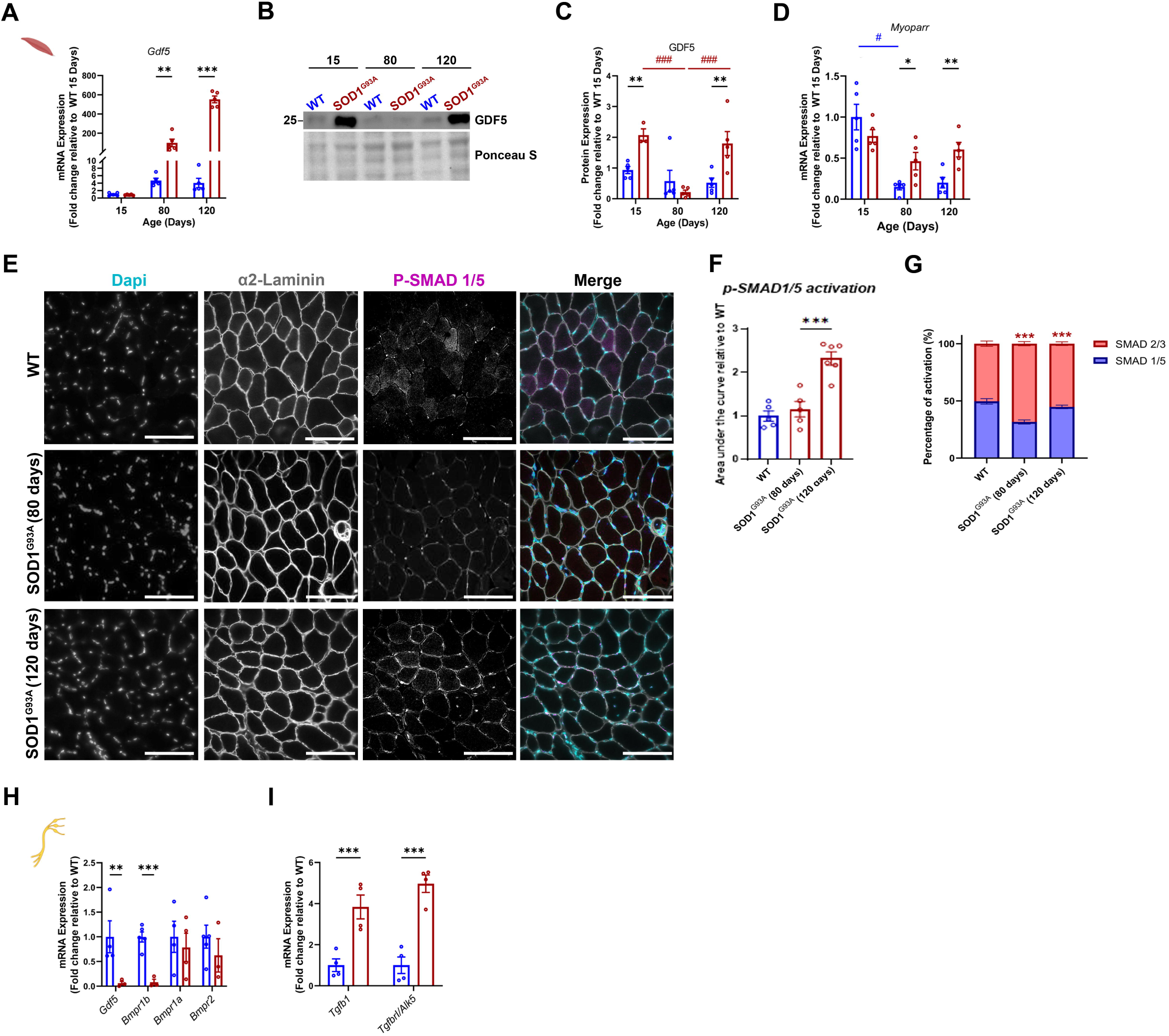
Impairment of the GDF5-SMAD 1/5/8 signaling across the neuromuscular axis. **A:** RT-qPCR of *Gdf5* transcript expression in TA muscle from WT and SOD1^G93A^ mice at P15, P80 and P120 (N=5) B: Representative Western blot of GDF5 protein levels in TA muscle at P15, P80 and P120 C: Quantification of Western blot GDF5 protein expression (fold change relative to WT at P15) in TA muscle at P15, P80 and P120 (N=5) D: RT-qPCR of the lncRNA *Myoparr* expression (fold change relative to WT at P15) in TA muscle from WT and SOD1^G93A^ mice at P15, P80 and P120 (N=5) E: Representative image of P-SMAD 1/5 nuclear accumulation from WT and SOD1^G93A^ TA at P80 by immunostaining of nuclei (DAPI) (Cyan), α2laminin (Grey) and P-SMAD1/5 (Magenta) F: Quantification of P-SMAD1/5 nuclear accumulation at P80 (N=5) G: Quantification of SMAD signaling balance (P-SMAD1/5 vs P-SMAD2/3) at P80 and P120 (N=5-6) H-I : RT-qPCR analysis in sciatic nerve at mid symptomatic showing (H) expression of *Gdf5* and *Bmpr1a*, *Bmpr1b* and *bmpr2* and (I) expression of *Tgfb1* and *Tgfbr1/ Alk5 (*N=4-5) Data are presented as means ± s.e.m. P-values were calculated by Two-way ANOVA or mixed effects analysis followed by Fisher LSD (A, C, D, H, I, J) ; and unpaired t-test (F)

Collectively, these data revealed a complex temporal regulation of GDF5 expression during disease progression in SOD1^G93A^ muscle. Although GDF5 protein expression is already elevated at the asymptomatic stage, signaling remains inactive, indicating an early intracellular blockade. During the symptomatic phase (P80-P90), sustained *Myoparr* expression suppresses GDF5 translation, preventing the establishment of a compensatory GDF5 response. Only at the late stage (P120) does GDF5 signaling become activated, suggesting that the endogenous protective response mediated by GDF5 is delayed and ultimately insufficient to counteract disease progression.

This assumption was confirmed by the fact that the GDF5 pathway, leading to the SMAD1/5 complex phosphorylation (p-SMAD1/5) and its nuclear accumulation, was not activated at P80 in SOD1^G93A^ muscle (Fig 2E-F) while, in contrast, the TGF-β SMAD2/3 pathway was significantly upregulated (Fig S1C). Analysis of upstream TGF-β pathway activators revealed that whereas *Myostatin* gene expression decreased from P80 onwards, *Tgfb1* transcript levels significantly increased starting at P80 (Fig S1D). This symptomatic shift drove a strong upregulation of atrophy-related downstream targets, including the transcripts of SMAD2/3-dependent E3 ubiquitin ligase *Murf1, Musa1* and *Atrogin1* starting at P80 (Fig S1E). Consequently, this led to a SMAD balance that remained favorable to the pro-atrophic process at both P80 and P120 (Fig 2G).

To determine if this imbalance also occurs in the peripheral nervous system, we analyzed the sciatic nerve at mid symptomatic. We found a significant downregulation of *Gdf5* and its high affinity receptor *Bmpr1b* (Fig 2H), while *Tgfb1* and *Tgfbr1* transcripts encoding respectively for TGF-β1 and TGF-βR1 were significantly increased (Fig 2I), suggesting a shift toward SMAD2/3 pathway chronic activation and a deleterious environment.(28)

Overall, these findings indicated that the muscle transcriptional attempt to compensate denervation-induced muscle atrophy via GDF5 is hindered by a *Myoparr*-mediated translational blockage and demonstrated a profound disruption of the SMAD signaling balance across the neuromuscular system. The shift toward TGF-β signaling, together with failure of the endogenous GDF5 response, creates a pro-atrophic molecular environment. These results identify the restoration of the BMP/ TGF-β balance as a critical therapeutic target for adult ALS treatment.

### Transcriptomic profiling reveals widespread molecular alterations in muscle of terminal-stage SOD1^G93A^

To gain a comprehensive and unbiased understanding of the molecular crisis occurring within the muscle during the terminal stage of the disease and to identify how the previously observed BMP/TGF-β imbalance fits into broader pathological process, we performed a whole transcriptome analysis (RNAseq) on TA of SOD1^G93A^ mice at P120 compared to WT (Fig 3A). Differential expression analysis revealed 2579 dysregulated genes, of which 1357 were downregulated and 1222 upregulated in SOD1^G93A^ muscle relative to WT (Fig 3B). Gene Ontology (GO) enrichment analysis of biological processes indicated that upregulated genes were associated with muscle and cytoskeleton development, negative regulation of the translation and ubiquitin-dependent proteolysis. Downregulated genes were mainly linked to muscle contraction, mitochondrial processes and metabolism, reflecting perturbations in oxidative stress related pathways (Fig 3C-D).

**Figure 3:**
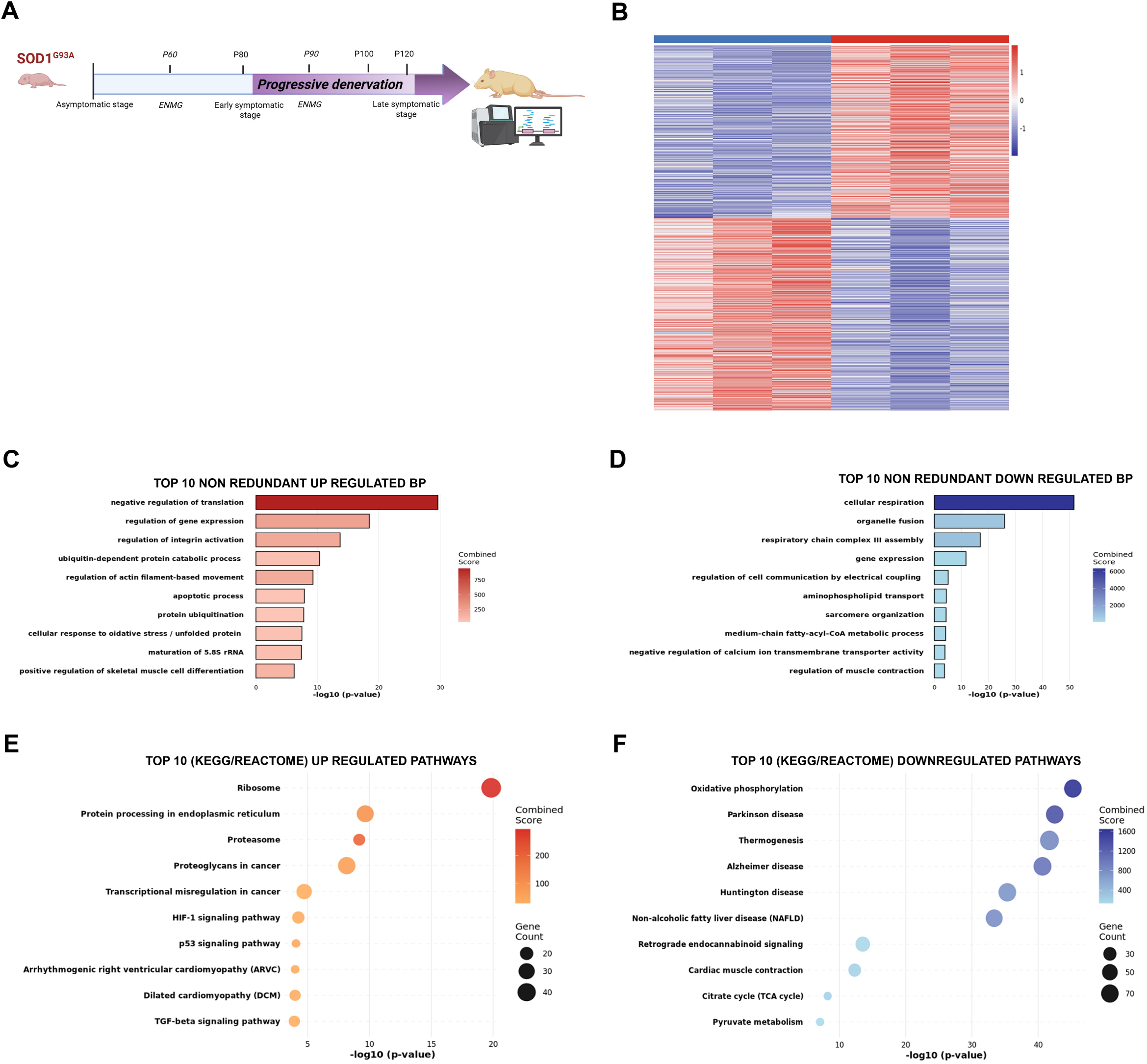
Transcriptomic profiling reveals widespread molecular alteration in muscle of terminal stage SOD1^G93A^. **A:** Schematic representation of the whole transcriptome analysis (RNAseq) experimental design on TA muscle from WT and SOD1^G93A^ mice at P120 (N=3) **B:** Heatmap depicting differential expression analysis with the 1222 upregulated and 1357 downregulated genes in SOD1^G93A^ muscle relative to WT **C-D:** Gene Ontology (GO) enrichment analysis of non-redundant biological process for (C) upregulated and (D) downregulated genes. **E-F:** Kyoto Encyclopedia of Genes and Genomes (KEGG) and Reactome pathways analysis for (E) upregulated and (F) downregulated pathways. Data are presented as Log2 fold change (B) and -log10 p-value (C-F). Differential expression analysis was performed using the DESeq2 package. P-values were calculated using the Wald test and adjusted by Benjamini-Hochberg test.

Integration of the Kyoto Encyclopedia of Genes and Genomes (KEGG), a knowledge base for systematic analysis of gene functions in terms of the networks of genes and molecule confirmed and extended these observations. Indeed, downregulated pathways consistently reflected mitochondrial and metabolic dysfunction as impaired oxidative phosphorylation and pyruvate metabolism. Upregulated pathway included apoptosis, ER stress, and, notably, TGF-β -SMAD2/3 signaling (Fig 3E, F).

Overall, transcriptomic profiling of SOD1^G93A^ muscles at terminal stage revealed a molecular signature dominated by a profound bioenergetic and proteostatic failures. This pathological state was coupled with the massive activation of the E3-ubiquitin ligase-mediated atrophic program. Furthermore, the global upregulation of the TGF-β signature at the transcriptomic level corroborates our previous biochemical observations (Fig 2), suggesting that the disruption of the BMP/TGF-β signaling balance could be a central driver of the progressive muscle decay.

Collectively, these data demonstrate that terminal SOD1^G93A^ muscle damages at terminal stage of the disease are governed by a global metabolic failure and a dominant pro-atrophic TGF-β environment. This molecular signature fully justifies targeting these signaling pathways to preserve muscle homeostasis during the course of the pathology.

### GDF5 OE in adult SOD1^G93A^ restores SMAD signaling balance and preserves muscle architecture

Our data established that the symptomatic stage of SOD1^G93A^ is characterized by the failure of the GDF5-mediated compensatory response to take hold in muscle and by a shift in the SMAD balance toward the pro-atrophic SMAD2/3 pathway. Given that GDF5 has been described as a potent neurotrophic and myotrophic factor capable of preserving muscle mass and neuromuscular integrity during aging(14,17), we investigated its therapeutic potential by overexpressing (OE) the GDF5 transgene in the SOD1^G93A^ at pre-symptomatic post-natal stage (P2) (Fig S2A-B). Analysis of functional parameters of these mice showed preservation of muscle mass at the terminal stage, normalization of CMAP at mid-symptomatic stage and extended survival in treated *SOD1^G93A^* mice compared to the un-treated animals (Fig S2).

Based on this initial proof-of-concept, we aimed to determine whether GDF5 could exert protective effects when administrated during a more clinically relevant window, specifically at the onset of the symptomatic phase. Thus, mice were injected by systemic route with AAV9-GDF5 (SOD1^G93A^-GDF5) or AAV9-Scramble (SOD1^G93A^-Scr) at P50 to allow time for transgene *GDF5* expression until the onset of symptoms at P80 (Fig 4A). Analyses were carried out at the end-point and we first confirmed GDF5 OE in both skeletal muscle and spinal cord by quantification of viral genome (Fig S3A-B) mRNA and protein for TA muscle (Fig S3A). To determine whether the GDF5 produced by the AAV9-GDF5 was functionally active and capable of counteracting the pathological TGF-β dominance observed in SOD1^G93A^ muscle, we assessed the activation status of the SMAD signaling balance. In one hand, we confirmed activation of the pathway by showing a significant increase of nuclear p-SMAD 1/5 labeling in SOD1^G93A^-GDF5 muscles compared to WT and SOD1^G93A^Scr (Fig 4B, C) and in another hand, immunostaining of p-SMAD2/3 showed a significant reduction of the proportion of positive nuclei (Fig S3C). These data provided evidence that GDF5 OE could inhibit the pro-atrophic TGF-β signaling and activate the trophic pathway in SOD1^G93A^ muscles (Fig 4D).

**Figure 4:**
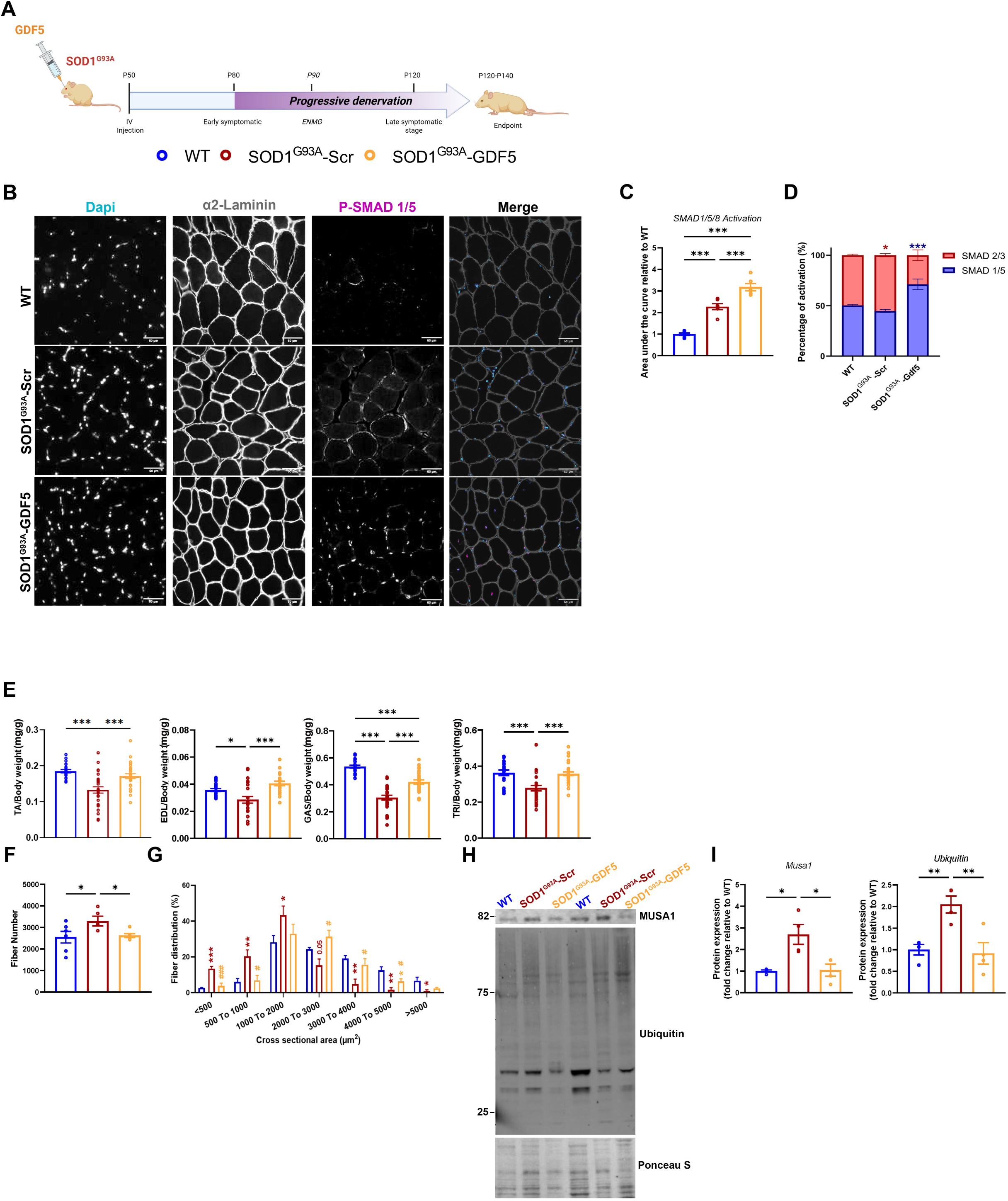
GDF5 OE in adult SOD1^G93A^ restores SMAD signaling balance and preserves muscle architecture. **A:** Schematic representation of the *in vivo* experimental design. Mice were systemically injected with AAV9-Scramble (SOD1^G93A^-scr) or AAV9-GDF5 (SOD1^G93A^-GDF5) at P50, ENMG were performed at P90 and other analyses were performed at the endpoint. **B:** Representative images of P-SMAD 1/5 nuclear accumulation from WT, SOD1^G93A^-scr and SOD1^G93A^-GDF5 TA at the endpoint by immunostaining of nuclei (DAPI), α2laminin and P-SMAD1/5 **C:** Quantification of P-SMAD1/5 nuclear accumulation from WT (Blue), SOD1^G93A^-scr (Red) and SOD1^G93A^-GDF5 (Orange) TA at the endpoint (N=6) **D:** Quantification of SMAD signaling balance (p-SMAD1/5 vs p-SMAD2/3) from WT, SOD1^G93A^-scr and SOD1^G93A^-GDF5 TA at the endpoint (N=6) **E:** Quantification of TA, Extensor digitorum longus (EDL), gastrocnemius (GAS) and triceps (TRI) muscle mass normalized on body weight at the endpoint (N= 20-28) **F:** Quantification of muscle fibers in whole TA cross-section at the endpoint (N=6) **G:** Distribution of fibers according to cross-sectional area at the endpoint (N=6) **H-I:** (H) Representative Western Blot and (I) quantification of MUSA1 protein levels and global protein ubiquitination profile in muscle lysates at the endpoint (N=4) Data are presented as means ± s.e.m. P-values were calculated by One-way ANOVA followed by Fisher LSD (C, F, I); Brown Forsythe and Welch ANOVA followed by Unpaired t with Welch’s correction (E, I), Kruskal-Wallis (E) and Two-way ANOVA followed by Fisher LSD (D and G).

The question was then to investigated how this GDF5 OE effect on SMAD molecular pathways could impact muscle mass homeostasis. Regarding morphometric analysis, GDF5 treatment led to a significant preservation of TA, Extensor longus digitorum (EDL) and triceps muscle mass of SOD1^G93A^-GDF5 compare to SOD1^G93A^Scr mice (Fig 4E). While SOD1^G93A^Scr muscles exhibited an increase of fiber number, likely due to fiber splitting, a constitutive hallmark of long-term denervation(29), this was normalized in SOD1^G93A^-GDF5 (Fig 4F). Indeed, fiber size distribution analysis revealed that GDF5 OE prevented the accumulation of small fibers (< 500; 500-1000 µm^2^) and promoted a distribution close to what observed for the WT muscles (Fig 4G). Importantly, given the known osteochondrogenic properties of the BMP family members, we verified that this GDF5 OE did not induce ectopic off-target effects. Alcian blue staining confirmed that this GDF5 induced mass and structural preservation occurred without any ectopic cartilage formation within the muscle tissue (Fig S3D) This structural preservation was associated to the down regulation of the transcript expression of the E3-ubiquitin ligases *Murf*1 and *Musa*1, two key downstream targets of the TGF-β pathway, as well as of the *atrogin-1*(11) (Fig S3E). Notably, the expression of MUSA1 was further validated at the protein level (Fig 4H, I). In addition, the global protein ubiquitination profile of SOD1^G93A^-GDF5 muscles was significantly restored to near WT level (Fig 4H, I).

Altogether, these findings establish that GDF5 OE acts as a proteostatic stabilizer by inhibiting the TGF-β driven catabolic program and acting as a myotrophic actor. GDF5 effectively prevents pathological fiber remodeling and maintains the structural and molecular integrity of the skeletal muscle despite the ongoing neurodegenerative process.

### GDF5 OE preserves NMJ integrity and restores neuromuscular transmission in adult SOD1^G93A^

As ALS is characterized by a progressive denervation leading to the structural functional collapse of the NMJ, it was essential to determine if benefits of GDF5 OE on muscle preservation could be extended to the synaptic compartment itself. We therefore investigated whether GDF5 OE in adult SOD1^G93A^ could prevent NMJ remodeling and rescue neuromuscular transmission throughout the disease progression by performing ENMG. We focused the functional analysis on the P90 timepoint which corresponds to the functional decline as shown in Fig1. ENMG analysis revealed a significant preservation of the CMAP amplitude in SOD1^G93A^-GDF5 compared to SOD1^G93A^-Scr (Fig 5A). To understand the molecular basis of this synaptic rescue, we analyzed markers of NMJ remodeling and synaptic support at the endpoint. We found that GDF5 treatment led to the normalization of the transcript expression of the embryonic AChR receptor subunits *Chrng (*gamma), particularly significant as its persistence is a hallmark of chronic denervation, and of the *Chrne (*epsilon) transcript (Fig 5B). Furthermore, we observed a significant increase in the transcript level of peri-synaptic SC specific markers, including *S100b* and *Mpz* (Fig 5B). Since peri-synaptic SC are essential for maintaining the synaptic cleft, their preservation suggests that GDF5 reinforces the glial support system at the NMJ.

**Figure 5:**
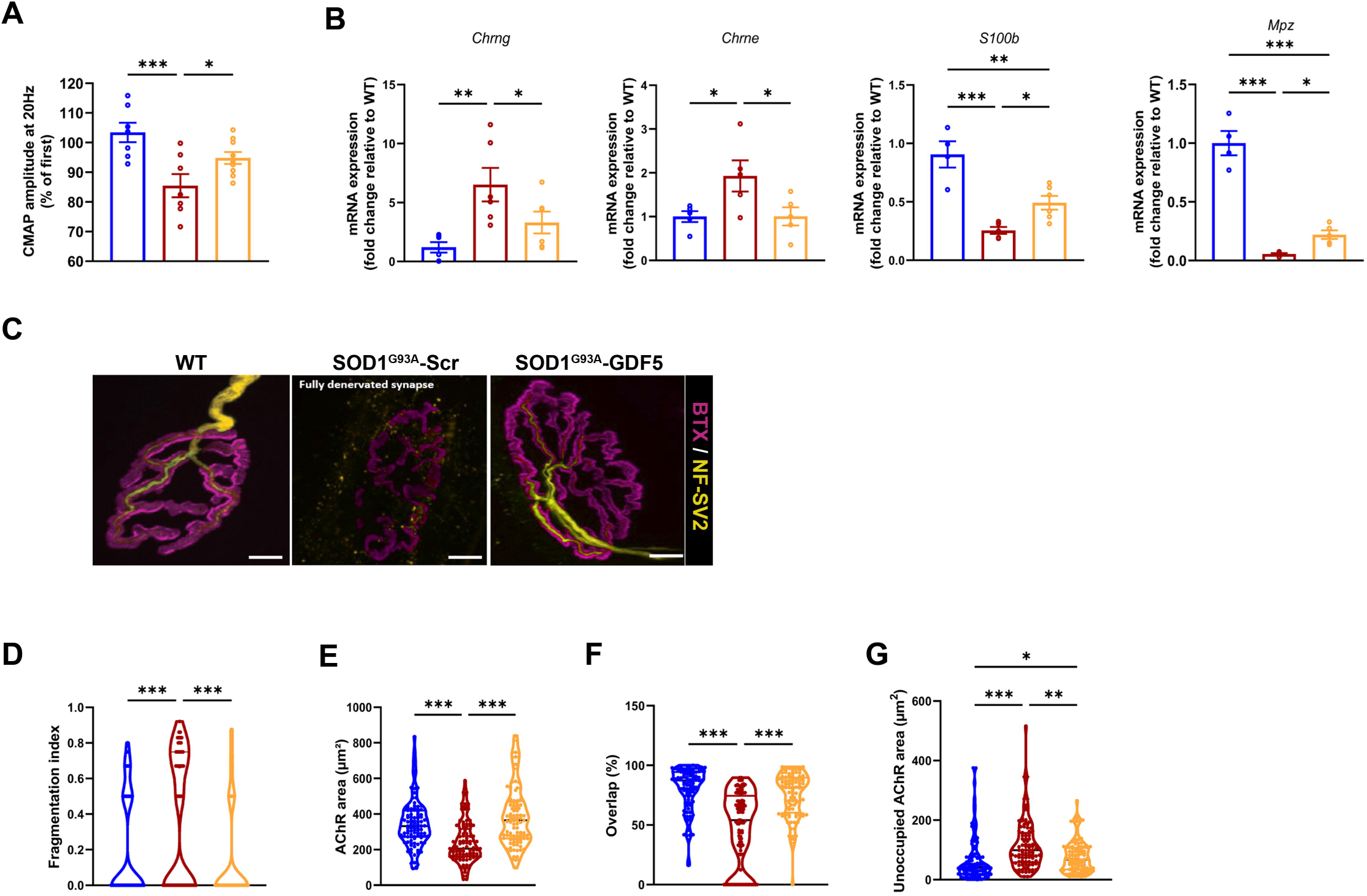
GDF5 OE preserves NMJ integrity and restores neuromuscular transmission in adult SOD1^G93A^. **A:** ENMG analysis at P90 quantifying the CMAP amplitude in WT (Blue), SOD1^G93A^-scr (Red) and SOD1^G93A^-GDF5 (Orange) TA (N=7-9) **B:** RT-qPCR of NMJ remodeling markers *Chrng*, *Chrne, S100b* and *Mpz* expression in TA at the endpoint (N=6) **C:** Representative images of NMJ morphology from WT, SOD1^G93A^-scr and SOD1^G93A^-GDF5 at the endpoint analyzed using NMJ morph. Presynaptic and post synaptic region are stained with NF200 +SV2 (Yellow) and α-bungarotoxin (Magenta), scale bar 10 µm **D-G:** Quantification of NMJ hallmarks including fragmentation index (D), total AChR area (E), unoccupied AChR area (F) and overlap percentage between pre- and post-synaptic component (G) (N= 100-102) Data are presented as means ± s.e.m. P-values were calculated by Ordinary One-Way ANOVA followed by Fisher LSD (A, B) and Kruskal-Wallis (D, E, F, G)

These functional and molecular benefits were corroborated by the morphological analysis of NMJs (Fig 5C). In SOD1^G93A^-GDF5 mice, we observed a stabilization of several key hallmarks of muscle denervation, including the fragmentation index and the AChR area (Fig 5D-E). GDF5 OE effectively restored the overlap percentage between pre- and postsynaptic regions (Fig 5F). Notably, while GDF5 OE significantly improved the unoccupied AChR area, this specific parameter was not fully rescued to WT levels (Fig 5G).

Overall, these data demonstrate that GDF5 OE acts as a potent synaptic stabilizer effectively sustaining the functional integrity of the NMJ throughout the disease progression by maintaining the glial niche, preserving postsynaptic excitability and preventing the structural fragmentation of the motor endplate.

### GDF5 OE counteracts ALS-associated oxidative stress and proteolysis signatures in muscle of adult SOD1^G93A^

Based on our findings demonstrating a terminal metabolic and proteostatic failure in muscles of SOD1^G93A^ mice (Fig 3), we sought to determine the extent to which GDF5 supplementation could reverse this pathological context. This question was addressed through a transcriptomic approach allowing a comparative analysis of SOD1^G93A^-Scr vs. SOD1^G93A^-GDF5 muscles (Fig 6A). Differential expression analysis identified 214 significantly regulated genes, including 83 upregulated and 131 downregulated transcripts (Fig 6B). Functional enrichment analyses revealed activation of pathways related to cellular component biogenesis, superoxide metabolism and BMP pathway, whereas pathways associated with proteolysis and negative regulation of reactive oxygen species (ROS) metabolism were significantly reduced in SOD1^G93A^-Scr compared with SOD1^G93A^-GDF5 muscles (Fig 6C,D). Consistent with these transcriptomic changes, GDF5 treatment enhanced ROS detoxification (Fig 6E) while reducing the expression of genes associated with protein degradation (Fig 6F). To validate these transcriptomic signatures and determine whether GDF5 directly affects muscle cells independently of neural degeneration, we employed a cell-autonomous model consisting of C2C12 myoblasts overexpressing human SOD1^G93A^ gene. Oxidative stress was measured using MitoSOX and confirmed that the C2C12-SOD1^G93A^ (Fig 6G) myotubes exhibited significantly higher mitochondrial ROS level compared to control C2C12. Continuous administration of recombinant GDF5 (rGDF5) throughout the differentiation process significantly reduced ROS accumulation. Furthermore, the pulse-chase protocol demonstrated that rGDF5 limited the initial generation of superoxide species when administered during early differentiation, while treatment restricted to the final 48 hours effectively reduced pre-existing ROS levels (Fig 6H). These *in vitro* findings confirm that mutant SOD1 expression is sufficient to induce cell-autonomous mitochondrial oxidative stress in muscle cells and demonstrate that GDF5 directly mitigates the intrinsic redox imbalance.

**Figure 6:**
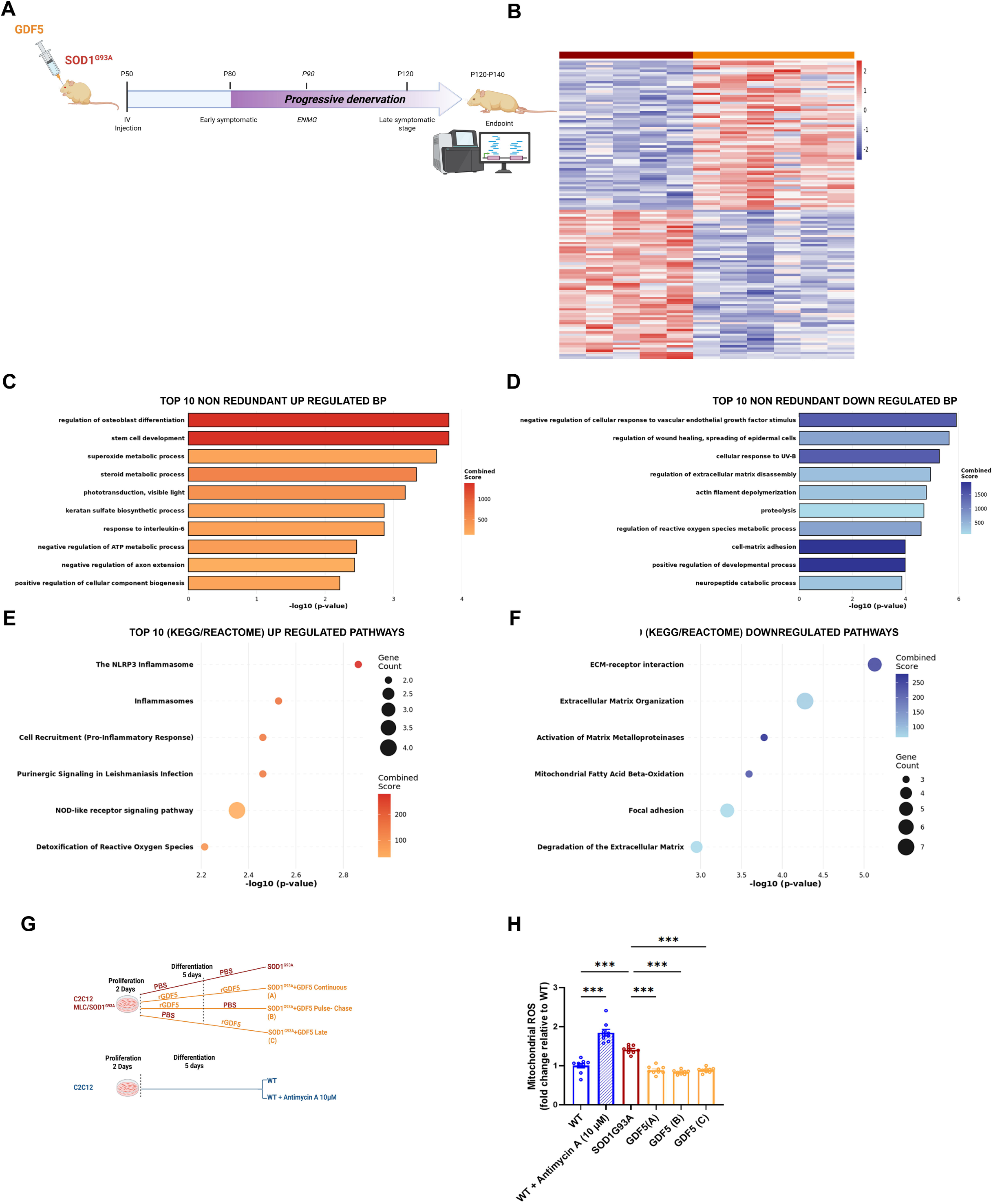
GDF5 OE counteracts ALS-associated oxidative stress and proteolysis signatures in muscle of adult SOD1^G93A^. **A** Whole transcriptome analysis (RNAseq) on TA muscle from SOD1^G93A^-scr and SOD1^G93A^-GDF5 mice at the endpoint (N=5-6) **B:** Heatmap depicting differential expression analysis with the 83 upregulated and 131 downregulated genes in SOD1^G93A^-GDF5 muscle relative to SOD1^G93A^-scr **C-D:** Gene Ontology (GO) enrichment analysis of non-redundant biological process for (C) upregulated and (D) downregulated genes **E-F:** Kyoto Encyclopedia of Genes and Genomes (KEGG) and Reactome pathways analysis for (E) upregulated and (F) downregulated pathways. **G:** Schematic representation of the *in vitro* experimental design using C2C12 WT and C2C12 MLC/hSOD1^G93A^ for mitoSOX oxidative stress assessment H: Quantification of mitochondrial ROS level (N=9) Data are presented as Log2 fold change (B), -log10 p-value (C-F) and means ± s.e.m (H). Differential expression analysis was performed using the DESeq2 package. P-values were calculated using the Wald test and adjusted by Benjamini-Hochberg test (B-F) and Brown Forsythe and Welch ANOVA test followed by Dunnett’s T3 test.

Collectively our results reveal that GDF5 OE acts as a multimodal regulator that neutralizes the toxic oxidative environment within the ALS motor unit. In addition, the ability of rGDF5 to reduce ROS accumulation in muscle cells expressing mutant SOD1 independently of neural input indicates that skeletal muscle contributes directly to disease-associated pathology and represents a therapeutically relevant target alongside neuroprotective strategies.

### GDF5 OE promotes molecular rehabilitation of motor neurons despite terminal neurodegeneration

Given the potent protective effect of GDF5 on the NMJ, we investigated whether GDF5 OE could extend this benefit to the central nervous system and improve global physiological outcomes. For this, we evaluated the impact of its overexpression on MN survival, the spinal inflammatory environment and the global survival of SOD1^G93A^-GDF5 mice. Despite significant preservation of the peripheral motor unit, GDF5 OE did not extend the overall survival of SOD1^G93A^ mice, as shown by Kaplan-Meier analysis (Fig 7A). Histological quantification of MN by ChAT staining in the cervical spinal cord revealed a significant and equivalent loss in both SOD1^G93A^-Scr and SOD1^G93A^-GDF5 groups compared to WT. Similarly, in the lumbar spinal cord, the trend toward MN preservation in GDF5 SOD1^G93A^ mice did not reach statistical significance at the terminal stage (Fig 7B-C). In striking contrast to this lack of structural rescue, GDF5 OE induced a substantial molecular change in the remaining MN. Indeed, we observed a remodeling of the spinal environment, characterized by a decrease in *Tgfb1* transcript level even lower that what observed for WT mice. This effect was accompanied by a normalization of the transcript level of vesicular acetylcholine transporter (*Vacht*) and *Apln* to WT level, alongside a marked upregulation of trophic and structural marker such as *Ngf* and neurofilament heavy chain *Nefh* transcripts in SOD1^G93A^-GDF5 compared to SOD1^G93A^-Scr mice (Fig 7D). Furthermore, GDF5 OE effectively reduced cellular stress, evidenced by the downregulation of *Gadd45a* and the restoration of *Gfap* mRNA levels (Fig 7D), suggesting that GDF5 limits the late-stage astrocytic degeneration typically observed in terminal stage of ALS disease in SOD1^G93A^mice.

**Figure 7:**
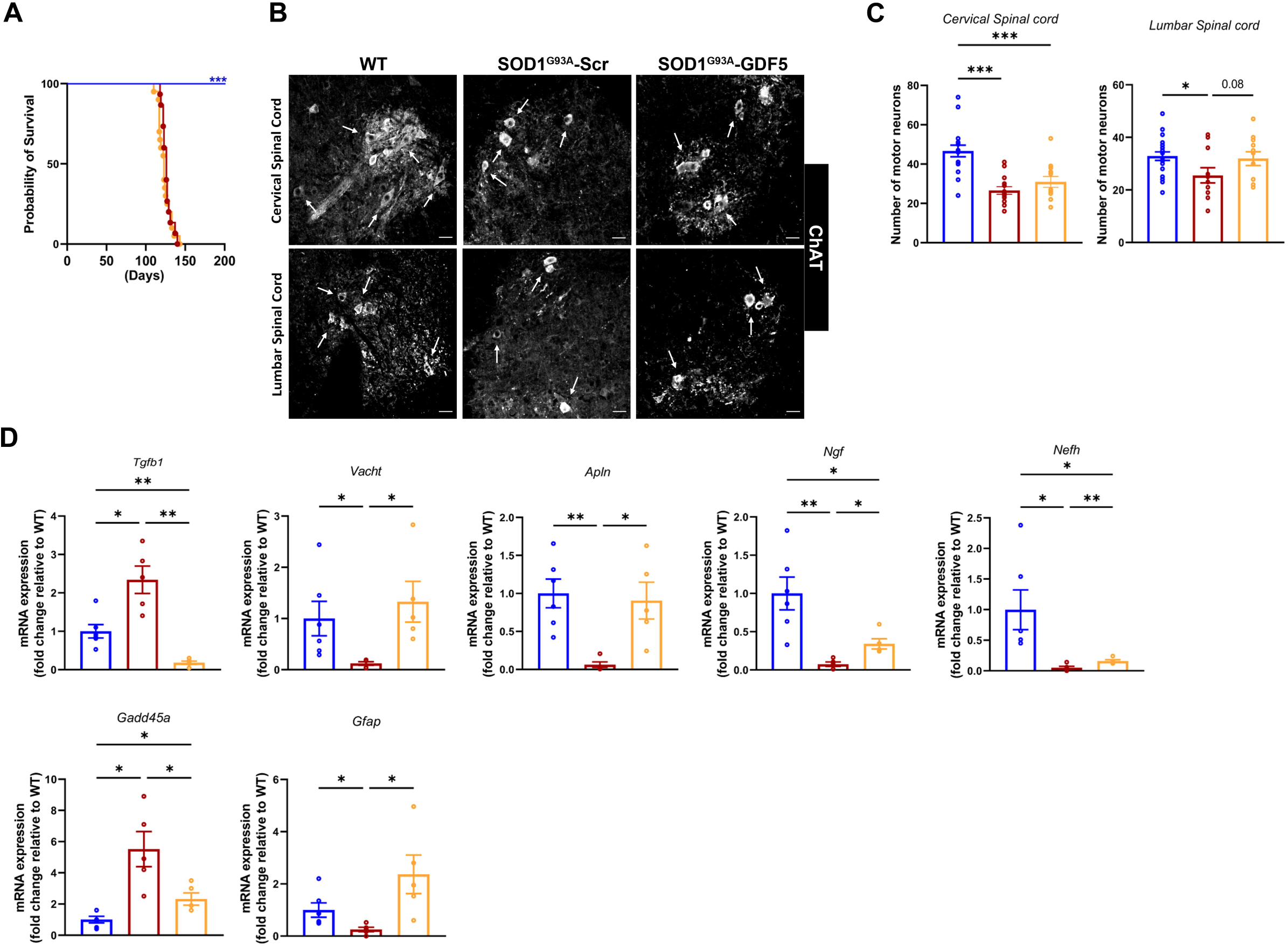
GDF5 OE promotes molecular rehabilitation of motor neurons despite terminal neurodegeneration. **A:** Kaplan-Meier survival analysis of WT, SOD1^G93A^-scr and SOD1^G93A^-GDF5 (N=15-20) **B:** Representative images of histological quantification of motor neurons by Choline acetyltransferase (ChAT) staining in the lumbar spinal cord **C:** Quantification of surviving motor neurons in the cervical and lumbar spinal cord (N= 11-20) **D-E**: RT-qPCR analysis of the spinal cord environment at the endpoint. (D) Expression of *Tgfb1, Vacht, Apln, Ngf* and *Nefh.* (E) Expression of stress and astrocytic markers *Gadd45a* and *Gfap* (N= 5-6) Data are presented as means ± s.e.m. P-values were calculated by Log-rank Mantel-Cox test (A), One-Way ANOVA followed by Fisher LSD (C), Brown Forsythe and Welch ANOVA followed by Unpaired t with Welch’s correction (C, D)

Taken together, these data demonstrate that while GDF5 OE fails to prevent terminal MN loss or extend lifespan in SOD1^G93A^ model, it significantly enhances the molecular homeostasis of the central motor system. This suggests that GDF5 influences molecular pathways associated with motor neuron maintenance, however, in the destructive context of the adult SOD1^G93A^, this molecular rescue remains insufficient to prevent the terminal phase of neurodegeneration or extend overall lifespan.

## Discussion

As a devastating neuromuscular disorder with no cure to date, ALS represents a critical public health challenge. Aiming to identify novel potential therapeutic targets to build clinical strategies significantly improving patient’s quality of life, our findings support the view that ALS involves dysfunction of the entire motor unit rather than a strictly neuron-centric neurodegenerative disease. We demonstrate that the progression of symptoms the SOD1^G93A^ model is not merely the consequence of motor neuron death, but an orchestrated collapse of the motor unit. By revealing the pathological upregulation of the lncRNA *Myoparr* in ALS, we highlight a molecular block that actively prevents skeletal muscle from mounting an effective compensatory response during the critical transition to the symptomatic phase. Bypassing this blockade through GDF5 OE, we showed a potent capacity for multimodal stabilization, where the muscle acts as an autonomous determinant of synaptic integrity and neuronal identity. Crucially, our findings established a biological dissociation between molecular rehabilitation and structural survival, suggesting that while the transcriptional fitness of the motor unit can be restored, terminal neurodegeneration remains gated by distinct central proteotoxic thresholds. Collectively, these findings support the exploration of therapeutic strategies targeting both central and peripheral components of the motor unit.

To capture this homeostatic breakdown across a precisely mapped chronological frame, we first characterized the progressive neuromuscular failure of the high-copy SOD1^G93A^ mouse model. Our initial physiological readout confirmed that symptomatic onset is defined by muscle atrophy and CMAP decrement before culminating in severe muscle wasting and NMJ fragmentation at the late symptomatic stage. Against this progressive pathological backdrop, a key discovery of our approach lies in the paradox identified at the pre-symptomatic stage, when SOD1^G93A^ muscles increase their expression of GDF5 protein despite both its mRNA level and *Myoparr* availability being unaffected. Because the transcription of *Gdf5* and its downstream targets remains strictly at baseline, the early post-transcriptional accumulation of GDF5 protein does not reflect an active response but rather a kinetic bottleneck in the ligand intracellular trafficking. This accumulation is biologically unproductive, as downstream target genes transcripts *Id1* and *Id2*(11) remain unexpressed. Given that conventional extracellular antagonists such as *Noggin*(30) and *Follistatin*(31) mRNA are not expressed at P15 in SOD^G93A^ mice, this suggests a strictly intracellular defect rather than extracellular sequestration of GDF5. Indeed, as a secreted member of the BMP family, GDF5 relies on folding within the endoplasmic reticulum (ER) and highly regulated transport through the Golgi apparatus(32). In ALS, ER stress is a well-established early pathological hallmark, which has been shown to begin as early as P14 in MN(33,34). It is essential to note that ER stress is known to disrupt the secretory pathway, inducing the intracellular retention of secreted proteins(35). These insights suggest that the translated GDF5 becomes trapped within a compromised secretory apparatus, preventing its normal release into the extracellular space. This defect in organelle proteostasis is further corroborated by our transcriptomic analysis at late disease stage, which highlights an enrichment in ER stress pathways in ALS muscles. Deprived of access to the extracellular space, GDF5 cannot bind to its receptor and its pathway remains functionally inactive.

This early secretory block eventually gives way to a dominant translational checkpoint during the symptomatic transition. At this stage, the lncRNA *Myoparr* instead of undergoing the physiological downregulation observed in WT tissues, maintains a sustained high expression level in SOD1^G93A^ muscles. By expanding the paradigm established by Hitachi et al to neurodegeneration, we show that this failure to downregulate *Myoparr* acts as a strict translational break, effectively halting GDF5 translation despite high mRNA availability(36). This blockade deprives the motor unit of its trophic shield, leaving it fully vulnerable to a pro-atrophic catabolic program. This symptomatic shift is driven by *Tgfb1* upregulation(10), a profound disruption in mitochondrial metabolism and the hyperactivation of downstream proteolytic pathways that characterize the end-stage ALS muscle, alongside the severe upregulation of atrophy-related E3 ubiquitin ligases(11). By overexpressing GDF5, we have been able to saturate and bypass this *Myoparr* blockade, demonstrating that muscle compensatory potential is merely suppressed rather than permanently lost during symptomatic progression.

Beyond its role in muscle proteostasis(11), our data establish GDF5 as a potent regulator of the neuromuscular interface, acting through both muscle-autonomous and retrograde mechanisms. Morphologically, GDF5 OE effectively arrested the structural dismantling of the motor endplate, as evidenced by the normalization of the fragmentation index and the restoration of synaptic overlap. This physical preservation is intrinsically linked to the stabilization of the tripartite synapse: GDF5 reinforced the peri-synaptic glial complex and prevented the pathological switch toward embryonic acetylcholine receptor subunit. Concurrently, our transcriptomic RNA-seq data at the endpoint confirm that the reactivation of the signaling pathway counteracted the catabolic program. This outcome is supported by the observation that GDF5 directly neutralized mitochondrial ROS generation and accumulation in C2C12-MLC/SOD1^G93A^ myotubes *in vitro*, redefining the muscle as an active participant in its own metabolic resilience.

The profound molecular rehabilitation of the peripheral motor unit holds particular therapeutic relevance when contrasted with the lack of systemic survival benefits observed in adult-treated mice. While GDF5 successfully improved the transcriptomic identity of spinal motor neurons, this retrograde support failed to arrest terminal death. This limitation likely stem from a dual pathological constraint. On one hand, the overwhelming central proteotoxicity inherent to the high-copy transgene(37), on the other hand, the accumulation of irreversible neurodegenerative damage prior to adult intervention in this model. This biological dissociation highlights a critical dual axis therapeutic requirement: while central-acting disease-modifying agents remain necessary to counteract cell-autonomous proteotoxicity within the spinal cord, targeting and stabilizing the motor unit at the periphery represents an equally indispensable mutation-independent strategy. Looking ahead, the clinical translation of this peripheral strategy is strongly supported by the therapeutic potential of recombinant GDF5. Indeed, we previously demonstrated that systemic administration of recombinant GDF5 successfully replicates these NMJ protective outcomes. This positions the recombinant protein as a highly viable, non-viral alternative. By focusing its action strictly on the peripheral component of the motor unit, this approach bypasses the safety concerns of viral gene therapy, offering a practical and scalable asset for future synergistic treatments in ALS. The fact that shifting the therapeutic window to an early postnatal stage successfully extends overall lifespan, demonstrates that early peripheral support can partially counteract pre-symptomatic degenerative events before irreversible structural damage occurs. Crucially, in this severe model, substantial lifespan extensions are typically achieved only by directly silencing the mutant *SOD1* transgene(7,38). However, neutralizing the genetic driver of this specific model can obscure downstream, mutation-independent mechanisms. By avoiding direct SOD1 silencing, our approach aimed to target the peripheral neuromuscular dismantling that universally characterizes human ALS pathology(39). The fact that early GDF5 supplementation succeeds in prolonging survival and preserving the integrity of motor unit underscores the broad therapeutic potential of this strategy. Indeed, since GDF5-mediated stabilization of the NMJ is independent of correcting a specific genetic defect, this approach could be effectively applied to all forms of ALS including genetic and non-genetic one.

In conclusion, our work supports a major paradigm shift, redefining ALS from a strictly cell-autonomous condition to a systemic pathology. This conceptual evolution paves the way for future synergistic therapeutic strategies, combining central-acting agents to halt neurodegeneration with peripheral stabilizer, such as GDF5 to maintain muscle function and protect the neuromuscular synapse in all ALS forms.

## Material and methods

### Animals

All animal procedures were reviewed and approved by an external ethics committee and the French Ministry of Higher Education and Scientific Research (Project authorization # 32044). Experiments were conducted in SOD1G93A mice, B6SJL-Tg (SOD1*G93A)1Gur/J (JACKSON SN 2726), were purchased from Jackson Laboratory. The animals were housed under SPF conditions under 12 h/12h light/dark conditions, with ad libitum access to food and water and cared for following the Directive 2010/63/EU. Endpoint were determined as the time when the mouse was unable to right itself in 30 seconds when placed on its side.

### Plasmids and AAV production

pSMD2-GDF5 have been generated by direct cloning of Gdf5 ORF (NM_008109.2), flanked by EcoRI and NheI sites (GeneArt string; ThermoFisher Scientific), in pSMD2 AAV2/9 (AAV9) under CMV promoter. pSMD2-Scramble have been generated by direct cloning of a scramble nucleotide sequence in pSMD2 and, under CMV promoter. AAV9 pseudotyped vectors have been prepared by the AAV production facility of the Center of Research in Myology, by transfection in 293 cells as described previously(14). The final viral preparations were kept in PBS solution at −80°C. The particle titer (number of viral genomes per ml) was determined by quantitative PCR.

### AAV Injections

For intravenous administrations, SOD1^G93A^ mice were injected at either 2 (P2) or 50 (P50) days of age. Newborn mice received a temporal vein injection of 70 µl of viral vector solution at a dose of 4.38 x 10^13^ vg/kg using using a Hamilton syringe (32G). Adult mice were anesthetized by an intraperitoneal injection of a ketamine/xylazine mixture (100 mg/kg ketamine and 10 mg/kg xylazine; 0.1 mL per 20 g of body weight). A total of 4,3810^13^ vg/kg of viral solution was administered into the retro-bulbar sinus using an insulin syringe (29G, Terumo) in a total of 50µL.

### Cell culture

C2C12 and C2C12 MLC/hSOD1^G93A^ cells were maintained in Dulbecco’s modified Eagle’s medium with 4.5g/L glucose (DMEM + GlutaMAX) supplemented with 20 % of fetal bovine serum, 1% Pen Strep (Gibco) and 2,5 μg/mL of puromycin (Biotechne) for C2C12 MLC/SOD^1G93A^ cells. To induce myogenic differentiation, cells were shifted to differentiation medium (DM), DMEM with 2% horse serum (Gibco) and 1% Pen Strep (Gibco).

### GDF5 treatment and mitoSOX assay

To evaluate the impact of GDF5 on oxidative stress, cells were treated with 200 ng/mL of recombinant mouse GDF5 (rGDF5 Sigma Aldrich) across three distinct temporal windows during differentiation: Continuous treatment from day 1 to day 5, Early treatment (Pulse-chase) from day 1 to day 3, Late treatment (Late) from day 3 to day 5.

Mitochondrial ROS production was assessed at day5 using mitoSOX green superoxide indicator (Thermofisher Scientific). Briefly, cells were rinsed with Hanks Balanced Salted solution without phenol red (HBSS Gibco) and incubated in the dark for 20 minutes at 37°C with 5 µM MitoSOX green. Antimycin A 10 µM (Sigma Aldrich) was used as a positive control for ROS induction. Fluorescence intensity was measured using a Tecan Spark microplate reader (Ex 490 nm /Em 520 nm). To account for variation in cell density, oxidative stress levels were normalized to total cell nuclei using DAPI staining fluorescence intensity (Ex 340 nm / Em 460 nm).

### Electroneuromyography measurements

Electroneuromyography (ENMG) measurements was achieved in TA muscle as previously described(14). During ENMG experiment mouse body temperature was maintained at 37°C with a heating plate. The sciatic nerve was stimulated with series of 10 stimuli at 20 Hz. Compound muscle action potentials (CMAP) were amplified (BioAmp, ADInstruments), acquired with a sampling rate of 100 kHz, filtered with a 5 kHz low-pass and a 1 Hz high-pass filter (Powerlab 8/25, ADInstruments). Peak-to-peak amplitudes were analyzed with LabChart 8 software (ADInstruments). In situ muscle force and ENMG experiments were performed in a blinded manner.

### Immunofluorescence and Histology

Animals were anesthetized by intraperitoneal injection of ketamine (100 mg/kg) and xylazine (10 mg/kg) at the end point and transcardically perfused with PBS. Tibialis anterior (TA), Gastrocnemius (GAS) and triceps (TRI) were mounted on tragacanth and frozen in cold isopentane. Extensor longus digitorum (EDL) were fixed in paraformaldehyde (PFA) 4% and sciatic nerve were collected in cryotube and flash frozen in nitrogen. Mice were then perfused with 4% paraformaldehyde (PFA) (Sigma-Aldrich) in PBS. The spines were explanted and stored in 4% PFA at + 4°C for at least 24 hr, then transferred into a PBS-sucrose solution (30%) and stored for at least 24 hr at 4°C. Spinal cords were extracted, embedded in Tissue-Tek (OCT), and frozen in cold isopentane. 14-μm-thick sections were serially cut from the whole spinal cord.

Muscle sections (12 μm thick) were performed on a cryostat (Leica Biosystems), fixed on glass slides and stored at -80°C. For Alcian blue staining, sections were fixed in 4% paraformaldehyde (PFA-Sigma-Aldrich) for 10 min, washed in PBS and then stained in Alcian Blue 8GX (Sigma-Aldrich) dissolved in 3% Acetic acid for 30 min and counterstain in Nuclear fast red 0.1% (Sigma-Aldrich) for 5 min. The muscle sections were further dried in gradually increasing concentration of ethanol/water solutions and, after fixation in 100% xylene, were mounted in Vectamount (Vector Laboratories).

For immunofluorescence procedures, muscle cryosections were rehydrated in phosphate-buffered saline (PBS), fixed or not with PFA 4% for 10 min, permeabilized with 0.5% Triton X-100 (Sigma-Aldrich) and blocked in a blocking buffer PBS containing 5% bovine serum albumin (BSA), 10% horse serum and 0.2% Triton X-100 for 1 h. Sections were incubated in blocking buffer with primary antibodies overnight at 4°C, washed in PBS, incubated for 1 h with secondary antibodies, thoroughly washed in PBS, incubated with 4′,6′-diamidino-2-phenylindole (DAPI) for nuclear staining for 10 min and mounted in Fluoromount (Southern Biotech). Spinal cord sections were also subjected to antigen retrieval with citrate buffer (10 mM citric acid, pH 6) at 85°C. Sections were blocked in a blocking buffer PBS containing 5% BSA, 10% normal donkey serum and 0.1% Triton X-100 for 1h. Sections were incubated in blocking buffer with a primary antibodies overnight at 4°C, washed in PBS, incubated for 1 h with secondary antibodies, thoroughly washed in PBS, incubated with DAPI for nuclear staining for 10 min and mounted in Fluoromount (Southern Biotech)

Imaging was performed on a confocal Nikon Ti2 microscope equipped with a motorized stage. To assess P-SMAD (1/5 and 2/3) activation, nuclear translocation was evaluated using a custom automated macro. The resulting fluorescence intensity values within the nuclei were fitted to a normal distribution, and the area under the curve was calculated to determine the proportion of P-SMAD-positive activated nuclei.

### Morphometric analysis

For fiber and nucleus detection, we used different artificial intelligence deep learning pretrained models. Fibers were segmented with Cellpose v3 using a model trained for cytoplasm. Nuclei were segmented using StarDist. The segmented image masks were then imported into QuPath, where we measured morphological features such as area and shape, and identified cells using the pixel intensity thresholding function. Finally, all measurements were exported to Excel, where we organized the data and performed statistical analyses such as CSA, fiber type classification, and staining positivity using thresholds as previously described.

### Immunoblotting

Cryosections from frozen TA muscles were homogenized with a dounce homogenizer in a cell lysis buffer (Cell Signaling) supplemented with phosphatase inhibitor cocktail (Roche). Lysates were centrifuged for 5 min at 1500g and protein concentration was determined in supernatant with Bradford method using Protein Assay Dye Reagent (Bio-Rad). Proteins were denatured at 95°C for 5 min with Laemmli buffer and β-mercaptoethanol (10% v/v) and then separated by electrophoresis (Nu-PAGE 4–12% Bis-Tris gel; Thermofisher Scientific) and transferred to nitrocellulose membranes (GE Healthcare). Membranes were blocked with 5% skimmed milk diluted in Tris Buffered Saline (TBS)-0.1% Tween (TBS-T) and incubated at 4°C overnight with primary antibodies. After washes in TBS-T, membranes were incubated with secondary antibodies conjugated to a fluorophore (Bio-Rad). Images were acquired with ChemiDoc™ MP (Bio-Rad) and band intensity was quantified using Image Lab software (Bio-Rad). Antibodies used are listed in table S2.

### Gene expression analysis

Total RNA was isolated from mouse muscles using TRizol (Life Technologies) and Direct-zol RNA Miniprep (Zymo research). From PFA-fixed spinal cord using RNeasy FFPE Kit (Qiagen) according to manufacturer protocol.

Complementary DNA was generated with Superscript II Reverse transcriptase (Life Technologies), and analyzed by real[time qPCR. Real-time qPCR were performed on QuantStudio 3 and 7 Pro Real-Time PCR System (Applied Biosystems) using Power SybrGreen PCR MasterMix (Applied Biosystems). All data were analysed using the 2-ΔΔCT method and normalized to P0 (mouse acidic ribosomal phosphoprotein P0) mRNA expression. The sample reference to calculate mRNA fold change is indicated in each panel. Primers used are listed in table S1.

### RNA-sequencing and Bioinformatic analysis

#### Library preparation and sequencing

RNA libraries were prepared using AccuraCode® technology (Singleron biotechnologies) according to manufacturer’s instructions. Muscles RNA samples were deposited in individual wells containing specific oligonucleotides harboring an illumina sequencing primer, a unique well barcode, a unique molecular identifier and a polyT sequence for mRNA capture. Following capture, cDNA was synthesized via a single-step RT-PCR. The resulting cDNAs were pooled, randomly fragmented and ligated with adapters to generate libraries compatible with Illumina platforms.

#### Data processing and Read alignment

After sequencing, the quality of raw data was assessed by filtering out low-quality of reads and monitoring error rates and GC content. High quality reads were aligned to Mouse mm10 (Ensembl) reference using the Spliced transcripts alignment to a reference (STAR) tool. Transcript quantification was subsequently perform using featureCounts.

#### Differential gene expression and enrichment analysis

For differential expression analysis, count data were normalized using the Variance Stabilizing Transformation (VST) within the DESeq2 package. For the WT versus SOD1^G93A^ comparison, differentially expressed genes (DEGs) were filtered as those with an adjusted p-value (padj) ≤0.05 (Benjamini-Hochberg method) and an absolute log2FoldChange (|log2FC|) ≥0.25. However, to account for the high inter-individual biological variability inherent to end point SOD1^G93A^ mice, in the SOD1^G93A^-Scr vs. SOD1^G93A^-GDF5 group, a relaxed threshold pf padj≤0.15 was applied for exploratory pathway screening. Functional enrichment analyses, including Gene Ontology (GO) and KEGG/Reactome pathways, were conducted using the R package enrichR. Reductions of GO redundant terms were conducted with the R package rrvgo. Crucially, all candidate target genes and downstream signalling pathway identified through this screening were systematically validated using independent, orthogonal methods, including RT-qPCR, western blotting and *in vitro* functional assays.

#### Viral genome quantification

Genomic DNA was extracted for mouse muscles, sciatic nerves and spinal cord with the PureGene tissue core Kit (Qiagen). AAV genome copy number and genomic DNA were quantified on 100ng of genomic DNA by real-time duplex qPCR. Real-time duplex qPCR were performed on QuantStudio 3 (Applied Biosystems). Primers were selected for specific amplification of the transgene promoter sequence for the viral vector and normalised on the quantity per nucleus. All genomic DNA samples were analysed in duplicate.

#### Neuromuscular-junction morphology

EDL muscles were immediately fixed 1 hour in 4% PFA after mice euthanasia. After washes in PBS, muscle fibers were isolated by mechanical dissociation. Isolated fibers were incubated 1 hour with 0.1M Glycine-PBS, then blocked in 4% BSA, 5% goat serum, 0.5% Triton-PBS and incubated at 4°C overnight with Neurofilament antibody diluted in blocking solution. Isolated fibers were washed several times in 0.1% Triton-PBS, then incubated at 4°C overnight with a secondary antibody (Table S3) and α-Bungarotoxin-Alexa Fluor™ conjugated 594. Isolated fibers were then mounted on glass slide with VECTASHIELD® Antifade Mounting Medium (Eurobio Scientific) and kept at 4°C until image acquisition. Images were acquired with a confocal microscope (Zeiss 63X objective) and edited with Zeiss Zen Lite 3.7 software. The same laser power and parameter setting were used throughout to ensure comparability. The images presented are single projected images obtained by overlaying sets of collected z-stacks. NMJ morphometric analysis was performed on confocal z-stack projections of individual NMJs by using ImageJ-based workflow adapted from NMJ-morph. At least 100 NMJs have been analysed per condition. NMJ morphometric analysis was performed in a blinded manner by the same investigator.

### Statistics analyses

For comparison between two groups, data were tested for normality using a Shapiro–Wilk test and for homoscedasticity using a Bartlett test followed by parametric (two-tailed paired, unpaired Student’s t-tests) or non-parametric test (Mann-Whitney) to calculate P values (as detailed in the figure legends). For comparison among more than two groups, according to normality ordinary one-way ANOVA or Kruskal-Wallis tests were performed. According to homoscedasticity, Brown-Forsythe ANOVA were performed. When it was necessary, two-way ANOVA tests were performed to compare between groups (as detailed in the figure legends). All ANOVA and Kruskal-Wallis were followed by appropriated post-hoc tests. All statistical analyses were performed with GraphPad Prism 9 software. Statistical significance was set at P < 0.05 and all bar graphs presented are means ± SEM. Asterisks indicate significant differences (*P < 0.05; **P < 0.01; ***P < 0.001) between groups, hashtags indicate significant differences (#P < 0.05; ##P < 0.01; ###P < 0.001) in same group between different timepoint according to statistical analysis performed.

## Author contributions

AB, MT, SF, PS and FPR conceived and designed the study. AB, SP, CG, TM, AF, MG, PM and JP performed the experiments. AB, SP and ZG analyzed the data. AB analyzed bioinformatic data. AB and FPR wrote the original draft. AM and GD writing review and editing. PS and FPR supervised the project.

## Acknowledgements

The authors acknowledge Association Institut de Myologie, Association Française contre les myopathies AFM-Téléthon to FPR.

The authors acknowledge the project M-BRAIN funded by the European Union – Next Generation EU – PNRR M6C2 – Investment 2.1 “Valorizzazione e potenziamento della ricerca biomedica del SSN to AM.

## Data availability statement

All data are available in the main text. Codes were used according to developer tutorial and available in the “Material and methods” section. The raw sequencing data of muscle RNAseq have been deposited in the National Center for Biotechnology Information (NCBI) in the Gene Expression Omnibus.

