## Supplemental Material for "GDF5 as a Multimodal Protector of the Motor Unit in Amyotrophic Lateral Sclerosis"

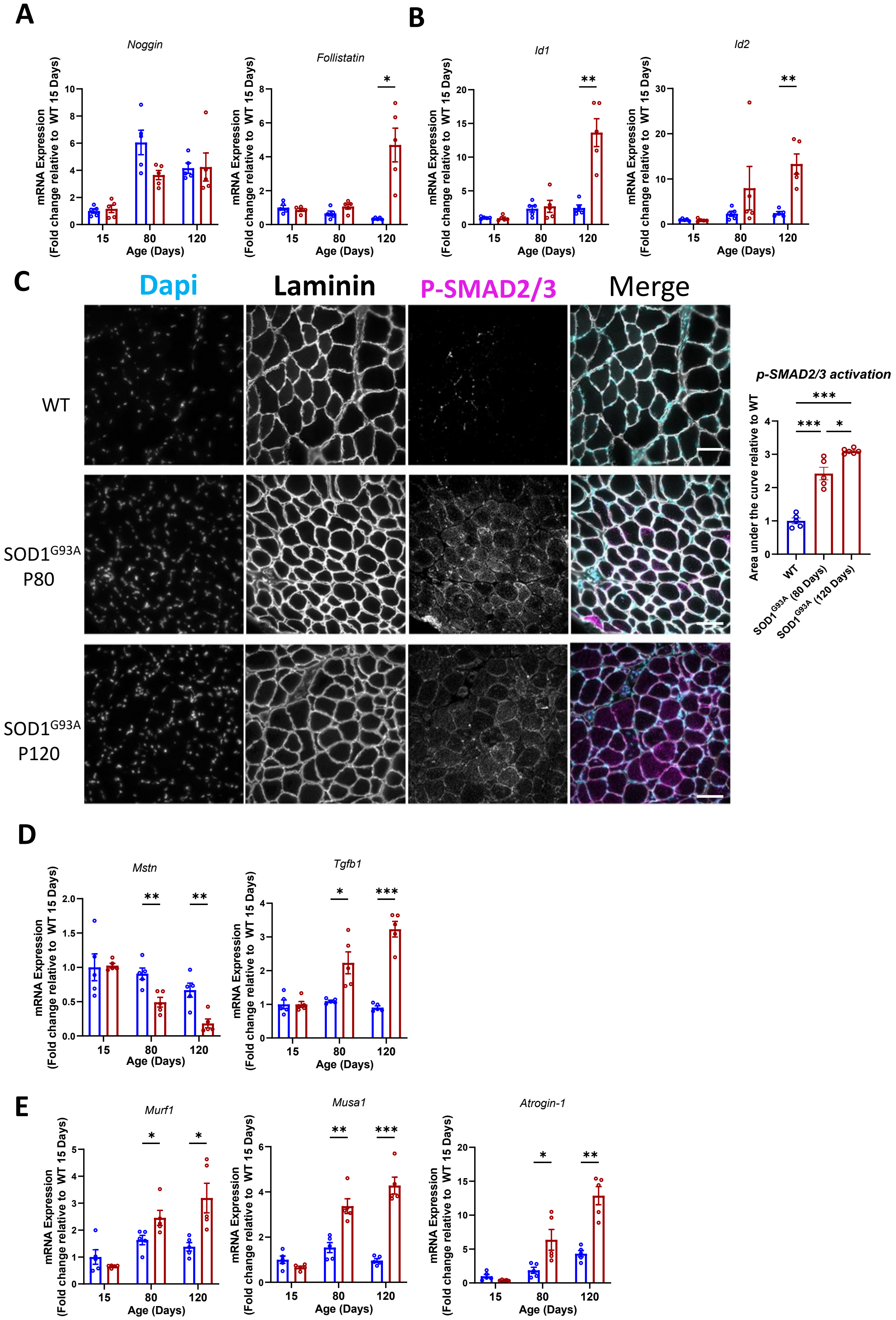

**Figure S1 : Characterization of BMP/TGF-β Ligands, antagonists and downstream target during the disease progression in SOD1$^{G93A}$ TA muscles**

**A:** RT-qPCR analysis of BMP/TGF-β inhibitor *Noggin* & *Follistatin* expression in TA muscle at P15, P80 and P120 (N= 5)
**B:** RT-qPCR analysis of BMP pathway downstream target *Id1* & *Id2* expression in TA muscle at P15, P80 and P120 (N= 5)
**C:** Representative image of pSMAD 1/5 nuclear accumulation and quantification from WT and SOD1$^{G93A}$ TA at P80 and P120 by immunostaining of nuclei (DAPI) (Cyan), α2laminin (Grey) and P-SMAD2/3 (Magenta)
**D:** RT-qPCR analysis of TGF-β pathway ligands *Mstn* & *Tgfb1* expression in TA muscle at P15, P80 and P120 (N= 5)
**E:** RT-qPCR analysis of TGF-β pathway downstream target *Murf1, Musa1* & *Atrogin-1* expression in TA muscle at P15, P80 and P120 (N= 5)
Data are presented as means ± s.e.m. P-values were calculated by Two-way ANOVA followed by a Fisher LSD (A,B, D,E) and Brown-Forsythe ANOVA followed by Unpaired t with Welch's correction (C).

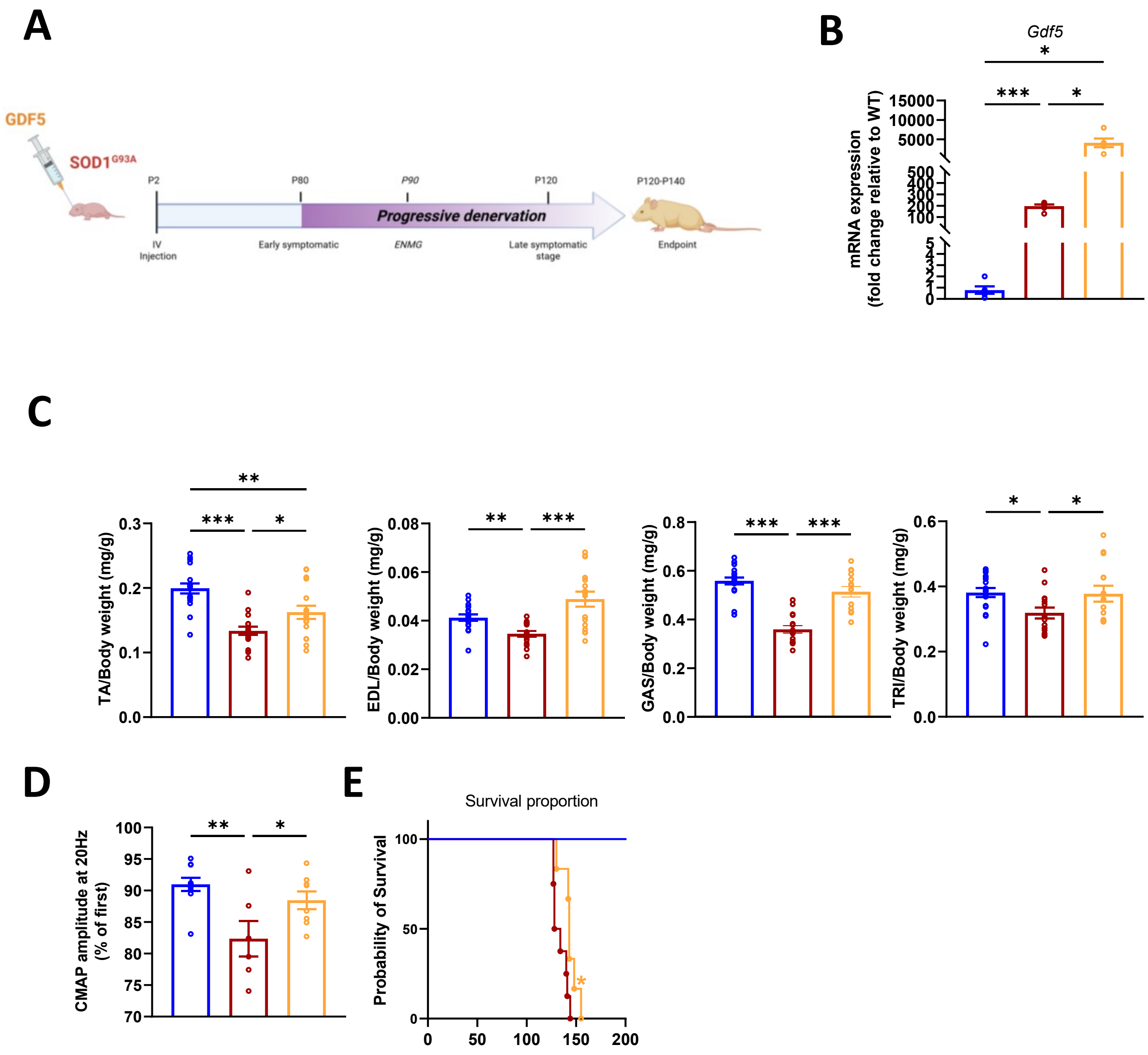


**Figure S2: GDF5 OE in newborn SOD1$^{G93A}$ mice improves muscle mass and function and enhances survival**

**A:** Schematic representation of the *in vivo* experimental design. Mice were systemically injected with AAV9-Scramble (SOD1$^{G93A}$-scr) or AAV9-GDF5 (SOD1$^{G93A}$-GDF5) at P2, ENMG were performed at P90 and other analyses were performed at the endpoint.
**B:** RT-qPCR of *Gdf5* transcript in TA muscles. (N=5-6)
**C:** Quantification of TA, Extensor digitorum longus (EDL), gastrocnemius (GAS) and triceps (TRI) muscle mass normalized on body weight at the endpoint (N=16-17
**D :** ENMG analysis at P90 quantifying the CMAP amplitude in WT (Blue), SOD1$^{G93A}$-scr (Red) and SOD1$^{G93A}$-GDF5 (Orange) TA (N=6-10)
**E:** Kaplan-Meier survival analysis of WT, SOD1$^{G93A}$-scr and SOD1$^{G93A}$-GDF5 (N=6-15)
Data are presented as means ± s.e.m. P-values were calculated by Brown-Forsythe ANOVA followed by Unpaired t with Welch's correction (B,C), One-Way ANOVA followed by Fisher LSD (D) and Mantel-Cox log Rank test (E)

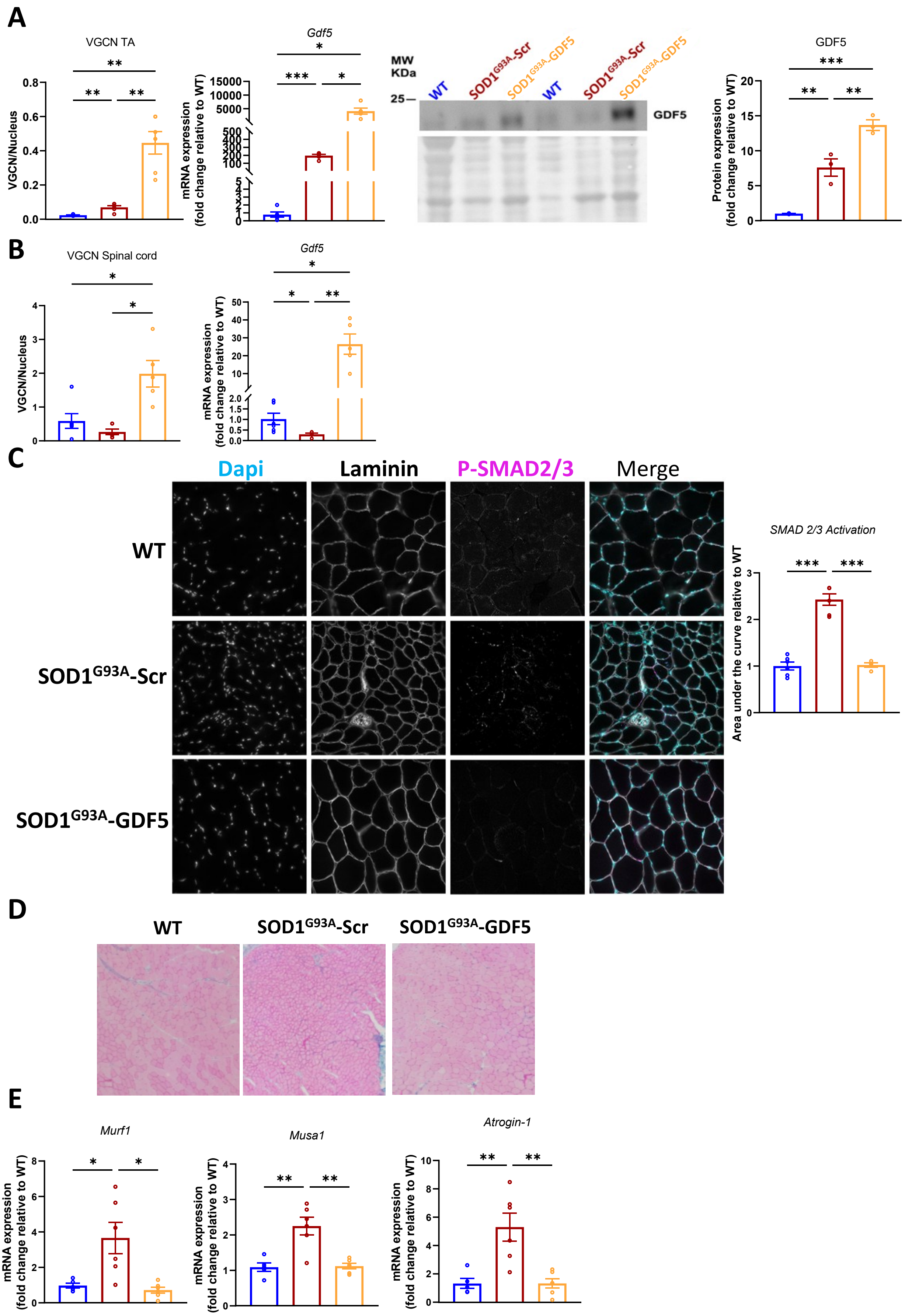
A
VGCN TA
VGCN/Nucleus
Gdf5
mRNA expression
(fold change relative to WT)
MW
KDa
25
WT
SOD1G93A-Scr
SOD1G93A-GDF5
GDF5
Protein expression
(fold change relative to WT)
B
VGCN Spinal cord
VGCN/Nucleus
Gdf5
mRNA expression
(fold change relative to WT)
C
Dapi
Laminin
P-SMAD2/3
Merge
WT
SOD1G93A-Scr
SOD1G93A-GDF5
SMAD 2/3 Activation
Area under the curve relative to WT
D
WT
SOD1G93A-Scr
SOD1G93A-GDF5
E
Murf1
Musa1
Atrogin-1
mRNA expression
(fold change relative to WT)

**Figure S3 : Validation of AAV-GDF5 biodistribution and signaling activation**

**A:** Quantification of Viral genome copy number, RT-qPCR of *Gdf5* transcript expression (N=6), Representative Western blot and quantification of GDF5 protein levels in TA muscles (N=3)
**B:** Quantification of Viral genome copy number and RT-qPCR of *Gdf5* transcript expression in Spinal Cord (N=5-6)
**C:** Representative image of P-SMAD 2/3 nuclear accumulation and quantification from WT and $SOD1^{G93A}$-scr and $SOD1^{G93A}$-GDF TA at the endpoint by immunostaining of nuclei (DAPI) (Cyan), α2laminin (Grey) and P-SMAD2/3 (Magenta) (N=6)
**D:** Representative image of Alcian Blue coloration
**E:** RT-qPCR analysis of TGF-β pathway downstream target *Murf1, Musa1* & *Atrogin-1* expression in TA muscle at P15, P80 and P120 (N= 6)
Data are presented as means ± s.e.m. P-values were calculated by Brown-Forsythe ANOVA followed by Unpaired t with Welch's correction (A,B,D), One-Way ANOVA followed by Fisher LSD (C)

**Listing of primers**

| Gene | Forward | Reverse |
|---|---|---|
| ***Chrna1*** | AAGCTACTGTGAGATCATCGTCAC | TGACGAAGTGGTAGGTGATGTCCA |
| ***Apln*** | CCTTGACTGCAGTTTGTGGA | CTCGAAGTTCTGGGCTTCAC |
| ***Atrogin1*** | GCCTTCAAAGGCCTCACG | CTGAGCACATGCAGGTCTGGG |
| ***Bmpr1A*** | GAAAGACCTGATTGACCAGTCC | CCCATCCATACTTCTCCATAGC |
| ***Bmpr1B*** | GAATACCAGCTTCCCTATCACG | TCTGCCTGAGACACTCATCACT |
| ***Bmpr2*** | TTGGACTCATCTACTGGGAGGT | TGGACACAAGAACCTGCATATC |
| ***Chrne*** | GCTGTGTGGATGCTGTGAAC | GCTGCCCAAAAACAGACATT |
| ***Chrng*** | GCTCAGCTGCAAGTTGATCTC | CCTCCTGCTCCATCTCTGTC |
| ***Folistatin*** | AGAGGAAATGTCTGCTTCCG | CACCTCTCTTCAGTCTCCTG |
| ***Gadd45A*** | GTGGTGTTGTGCCTGCTG | AGGATGTTGATGTCGTTCTCG |
| ***Gap43*** | CCTGCTGCTGTCACTGATGCTG | CCTGCTGCTGTCACTGATGCTG |
| ***Gdf5*** | ATGCTGACAGAAAGGGAGGTAA | GCACTGATGTCAAACACGTACC |
| ***Gfap*** | GAAGCTCCAAGATGAAACCAAC | TCCAGCGATTCAACCTTTCTC |
| ***Id1*** | AGTGAGCAAGGTGGAGATCC | GATCGTCGGCTGGAACAC |
| ***Id2*** | CTCCAAGCTCAAGGAACTGG | ATTCAGATGCCTGCAAGGAC |
| ***Lif*** | GTCTTGGCCGCAGGGATTG | GCACAGGTGGCATTTACAGG |
| ***Mpz*** | CCCTGGCCATTGTGGTTTAC | CCATTCACTGGACCAGAAGGAG |
| ***Mstn*** | AGTAAAAGCCCAACTGTGGATA | TCATGTCAAGTTTCAGAGATCG |
| ***Murf1*** | CGACCGAGTGCAGACGATCAT | GTGTCAAACTTCTGACTCAGC |
| ***Musa1*** | TCGTGGAATGGTAATCTTGC | CCTCCCGTTTCTCTATCACG |
| ***Musk*** | TTCAGCGGGACTGAGAAACT | TGTCTTCCACGCTCAGAATG |
| ***Nefh*** | TGCCGCTTACAGAAAGCTC | GCGTGGATATGGAGGGAATTT |
| ***Ngf*** | GCAGTGAGGTGCATAGCGTA | CTGTGTCAAGGGAATGCTGA |
| ***Noggin*** | GAAGTTACAGATGTGGCTGTGG | CACAGACTTGGATGGCTTACAC |
| ***S100β*** | CTTCCTGGAGGAAATCAAGGAG | CTCATGTTCAAAGAACTCATGGC |
| ***TGFb1*** | GGAGAGCCCTGGATACCAAC | GAAGTTGGCATGGTAGCCCTTG |
| ***Tgfbr1*** | TCCTCGAGATAGGCCGTTTG | GGCCAGGTGATGACTTTACAGTAG |
| ***VAChT*** | CCCTTTTGATGGCTGTG | GGGCTAGGGTACTCATTAGA |

**Listing of primary antibodies**

| Name /Target | Source | Reference | Species | Application | Dilution |
|---|---|---|---|---|---|
| GDF5 | Santa Cruz | sc-373744 | Mouse | WB | 1/500 |
| P-SMAD 1/5 | Cell signaling | 9516S | Rabbit | IF | 1/800 |
| P-SMAD 2/3 | Cell signaling | D27F4 | Rabbit | IF | 1/800 |
| MUSA1 | Santa Cruz | sc-514862 | Mouse | WB | 1/100 |
| Ubiquitin (K48-linkage Specific Polyubiquitin) | Cell signaling | 8081S | Rabbit | WB | 1/1000 |
| ChAT | Millipore | AB144P | Goat | IF | 1/100 |
| Neurofilament | Millipore | AB9568 | rabbit | IF | 1/500 |
| SV2-c | DSHB | | Mouse | IF | 1/500 |
| α Bungarotoxin-Alexa Fluor™ 594 | Thermofisher | B13423 | B. multicinctus | IF | 1/500 |
| Laminin α2 | Santa Cruz | sc-59854 | Rat | IF | 1/50 |
